# Homeotic Regulation of Olfactory Receptor Choice via NFI-dependent Heterochromatic Silencing and Genomic Compartmentalization

**DOI:** 10.1101/2020.08.30.274035

**Authors:** Elizaveta Bashkirova, Kevin Monahan, Christine E. Campbell, Jason M. Osinski, Longzhi Tan, Ira Schieren, Gilad Barnea, X. Sunnie Xie, Richard M. Gronostajski, Stavros Lomvardas

## Abstract

Expression of one out of >1000 olfactory receptor (OR) genes is stochastic but, yet, spatially organized in stereotypic anatomical segments, or “zones”, along the dorsoventral axis of the mouse olfactory epithelium. We discovered that zonal OR expression is specified by OR chromatin structure and genome architecture during olfactory neuron differentiation. Specifically, across every zone dorsally expressed ORs have higher levels of heterochromatic marks and long-range contacts than ORs expressed ventrally. However, OR heterochromatin levels and frequency of genomic contacts between ORs gradually increase towards ventral zones. Consequently, ORs from dorsal indexes accumulate high H3K9me3/H3K79me3 enrichment and become silenced in ventral zones, while ORs from ventral indexes lack activating long-range genomic interactions and, thus, cannot be chosen in dorsal segments. This process is regulated by NFIA, B, and X gradients along the dorsoventral axis, triple deletion of which causes homeotic transformations on zonal OR expression, heterochromatin formation, and genomic compartmentalization.

## Introduction

Pattern formation governs fundamental developmental processes in plants and animals, vertebrates and invertebrates. From the classic theoretical predictions on the chemical basis of morphogenesis(Turing, 1953), to the seminal demonstration of the genetic control of segmentation in drosophila(Nusslein-Volhard and Wieschaus, 1980); and from the elucidation of motor neuron pool identity in mammals (Dasen et al., 2005), to the explanation of photoreceptor patterning in invertebrate ommatidia(Johnston et al., 2011), the molecular and genetic underpinnings of developmental patterning are -at various extents-understood. However, the mechanisms that regulate developmental patterning in the mammalian olfactory system are completely unknown. In this sensory system, spatial patterns are apparent in the periphery, the main olfactory epithelium (MOE) and the olfactory bulb (OB)(Sullivan et al., 1995; Sullivan et al., 1994), but not in higher order brain processing centers(Stettler and Axel, 2009). In the MOE expression of >1000 class II olfactory receptor (OR) genes(Buck and Axel, 1991) from 18 different chromosomes(Sullivan et al., 1996; Zhang and Firestein, 2009) is organized in continuous “zones”, that are faithfully represented by zonally-restricted neuronal projections to corresponding areas of the OB(Ressler et al., 1994). Although there is evidence that the zonal organization of sensory inputs has behavioral and perceptual significance(Kobayakawa et al., 2007), without knowing how zonal organization is established we cannot evaluate the significance of zonal OR expression in olfactory perception.

Zones are stereotypic and evolutionarily conserved subdivisions of the MOE along the dorsoventral axis that express preferentially 50-200 OR genes from the whole OR repertoire. Originally, RNA ISH experiments revealed that the MOE is divided in 4 broad zones (Ressler et al., 1993; Vassar et al., 1993), with highly overlapping borders(Miyamichi et al., 2005). scRNA-seq experiments in micro-dissected MOE pieces assigned zonal coordinates to the majority of the ORs and redefined the anatomical subdivisions of the MOE into 5 continuous zones (Tan and Xie, 2018), with zone 1 being the most dorsal and zone 5 the most ventral. The septal organ, a distinct anatomical structure of the MOE expressing predominantly a single OR could be viewed as a distinct MOE zone(Tian and Ma, 2004), and so could the cul-de-sac MOE segments that express MS4A chemoreceptors(Greer et al., 2016). In fact, comprehensive 3D imaging of OR expression by multiplex RNA ISH, raised the prospect of 9 or more OR zones in the MOE(Zapiec and Mombaerts, 2020). Beyond these anatomical distinctions that are established along the dorsoventral axes of the MOE, additional olfactory sensory neuron (OSN) subdivisions exist that do not entail spatial segregation of molecularly distinct cell types. For example, OSNs expressing class I ORs (fish-like ORs)(Bozza et al., 2009), or trace amine associated receptors (TAARs)(Johnson et al., 2012), are committed to each of these chemoreceptor subfamilies but are intermingled with class II OR-expressing OSNs in their respective zones. Thus, the emerging picture suggests a hierarchical process that starts with broad zonal specification and commitment to a chemoreceptor family, followed by a singular stochastic choice of one of the zonally enabled receptor alleles. Although we made strides in the understanding of how this singular stochastic choice is accomplished, the molecular mechanisms that restricts it within a zonal repertoire are unknown.

The process of singular OR gene choice starts in OSN progenitors, where the already H3K9me2-positive OR genes, become marked with H3K9me3/H4K20me3, two hallmarks of constitutive heterochromatin (Magklara et al., 2011). Heterochromatin assembly is followed by the formation of OR-exclusive interchromosomal compartments (Armelin-Correa et al., 2014; Clowney et al., 2012; Tan et al., 2019) that further silence OR transcription by insulating OR genes from activating transcription factors(Clowney et al., 2012). OR compartments, which assemble due to downregulation of Lamin b receptor (Lbr) in OSN progenitors, also contribute to singular OR choice, by promoting assembly of an adjacent multi-chromosomal enhancer hub that associates with one of the previously silenced ORs, activating its transcription (Clowney et al., 2012; Markenscoff-Papadimitriou et al., 2014). This multi-enhancer hub consists of intergenic OR enhancers, the Greek Islands, which are bound by Lhx2, Ebf and Ldb1(Markenscoff-Papadimitriou et al., 2014; Monahan et al., 2019; Monahan et al., 2017). This tripartite protein complex promotes the highly specific interchromosomal contacts between Greek Islands, creating a strong transactivating complex responsible for the de-silencing and robust transcriptional activation of one of the compartmentalized OR alleles. Translation of the chosen OR in the ER induces the unfolded protein response pathway(Dalton et al., 2013) and generates a feedback signal(Lewcock and Reed, 2004; Serizawa et al., 2003; Shykind et al., 2004) that hinders the ability of the Greek Island hub to de-silence additional OR alleles, stabilizing singular OR gene choice for the life of the OSN(Lyons et al., 2013).

The aforementioned complexity in OR gene regulation likely compensates for the fact that traditional mechanisms of combinatorial gene regulation cannot determine which OR is activated in each OSN. Numerous lines of evidence support the notion that OR gene choice is a stochastic process. First, “allelic exclusion”, expression from only the maternal or paternal allele in each OSN, suggests that *cis* and *trans* regulatory information cannot sufficiently define which OR will be expressed in each cell(Chess et al., 1994). In this note, transgenic ORs, from minigenes(Vassalli et al., 2002) to YAC transgenes(Ebrahimi et al., 2000; Serizawa et al., 2000), are not co-expressed with the endogenous OR alleles from which they were derived, arguing against a deterministic mode of OR regulation. Second, OR gene switching, a low frequency switch from the original chosen OR allele to a different OR identity, further supports the OR multipotency of each OSN (Shykind et al., 2004). Third, scRNA-seq studies revealed that there is a transient differentiation stage where OSN progenitors co-transcribe seemingly random combinations of 10-15 OR genes(Hanchate et al., 2015; Saraiva et al., 2015; Tan et al., 2015). These results taken together suggests that an unknown mechanism integrates the position of the OSN with the identity of each OR to restrict this stochastic choice within a spatially determined OR gene combination.

To obtain insight at the remarkable problem of spatially determined randomness, we performed a comparative analysis of the transcriptome, chromatin modification landscape, and nuclear architecture of various cell populations from different MOE zones. Our data reveal that early OSN progenitors from every zone co-express predominantly zone 1 ORs. However, as we transition from zone 1 to zone 5 (i.e. from dorsal to ventral zones), developmental progression results in silencing of zone 1 ORs and activation of ORs expressed ventrally, culminating in the singular expression of ORs with the correct zonal identity. These spatiotemporal transcriptional transitions are regulated by two mechanistically linked processes: marking of an increasing number of OR genes with H3K9me3 and H3K79me3, and gradual recruitment of an increasing number of OR genes to OR compartments, which enables interactions with the Greek Island hub. Thus, within a zone, ORs with more dorsal identities have the highest levels of H3K9me3/H3K79me3 and most frequent long-range genomic contacts. However, because both heterochromatin formation and OR compartmentalization progressively increase towards ventral zones, in each zone there are three types of ORs: ORs expressed in previous (i.e. more dorsal) zones that have high levels of H3K9me3/H3K79me3 that keep them silent; ORs expressed in subsequent (i.e. more ventral) zones that lack both H3K9me3/H3K79me3 and activating long-range genomic interactions; and ORs with the correct zonal identity that have intermediate enrichment of H3K9me3/H3K79me3 and Hi-C contact frequencies. Thus, only ORs from the correct zonal index combine reversible levels of heterochromatin with the ability to interact with other ORs and with the Greek Island hub. This process is controlled by three paralogous transcription factors, NFI A, B, and X whose gradient expression in OSN progenitors increases towards zone 5, expanding both the number of ORs marked by H3K9me3/H3K79me3 and the zonal constitution of OR compartments. Triple NFI deletion in OSN progenitors transforms the heterochromatic landscape and the nuclear compartmentalization of zone 4/5 OSNs to patterns reminiscent of zone 2/3 OSNs, while it induces ectopic expression of zone 2/3 ORs in place of zone 4/5 ORs. In contrast triple NFI deletion in mature OSNs has no effect on zonal OR transcription. Thus, NFI genes act as homeotic regulators of OR gene choice that specify the zone of expression for each OR via chromatin-mediated silencing and genomic compartmentalization.

## Results

### OR transcription undergoes a zonal transformation during OSN differentiation

Before the onset of singular OR expression, mitotically active OSN progenitors co-express multiple OR genes at levels that are high enough for detection by scRNA-seq(Hanchate et al., 2015; Saraiva et al., 2015; Tan et al., 2015). To identify the developmental stage when zonal restrictions are established, we asked if the co-transcribed ORs follow zonal patterns of expression in defined populations of OSN progenitors. We used FACS to isolate specific OSN progenitor populations from Mash1CreER; tdTomato; Ngn1-GFP triple transgenic mice (Figure 1A). Mash1CreER is expressed in pluripotent stem cells of the MOE(GBCs)(Rodriguez-Gil et al., 2015), whereas Ngn1-GFP is predominantly expressed in immediate neuronal precursors (INPs)(Fletcher et al., 2017) and immature OSNs (iOSNs)(Magklara et al., 2011). This genetic strategy also allows isolation of newly differentiated mature OSNs (mOSNs), which recently turned off Ngn1-GFP transcription but still have detectable levels of GFP protein (Figure 1A). We injected P4 mice with tamoxifen, inducing permanent tdTomato expression, and then collected cells 48 hours later (Figure 1A). From each dissection we isolated four major cellular populations: tdTomato^+^/GFP^-^, tdTomato^+^/GFP^+^, tdTomato^-^/GFP^+bright^, tdTomato^-^/GFP^+dim^ (Figure 1A), which represent GBCs, INPs, iOSNs, and mOSNs respectively. First, we dissected the developing MOE into dorsal and ventral pieces, containing predominantly zones 1-2 and 3-5, respectively. scRNA-seq of these FAC-sorted populations confirmed that our genetic strategy enriches for the 4 major differentiation stages, with INP cells further divided into 3 subpopulations, INP1-3 (Supplemental Figure S1A). Moreover, tSNE plots verified in, an unbiased fashion, the developmental relationships of the 4 FAC-sorted cell populations (Figure 1B). OR mRNAs are first detected in INP3 cells (Figure 1C), which express multiple ORs, consistent with previously reported OR co-expression. Surprisingly, if we plot the zonal identity of the most highly expressed OR in each cell, we find that INP3 cells from both zonal dissections predominantly express zone 1 and 2 ORs. It is not until the OSN stage that cells from more ventral zones switch to expressing zone 3-5 ORs (Figure 1C).

**Figure 1:**
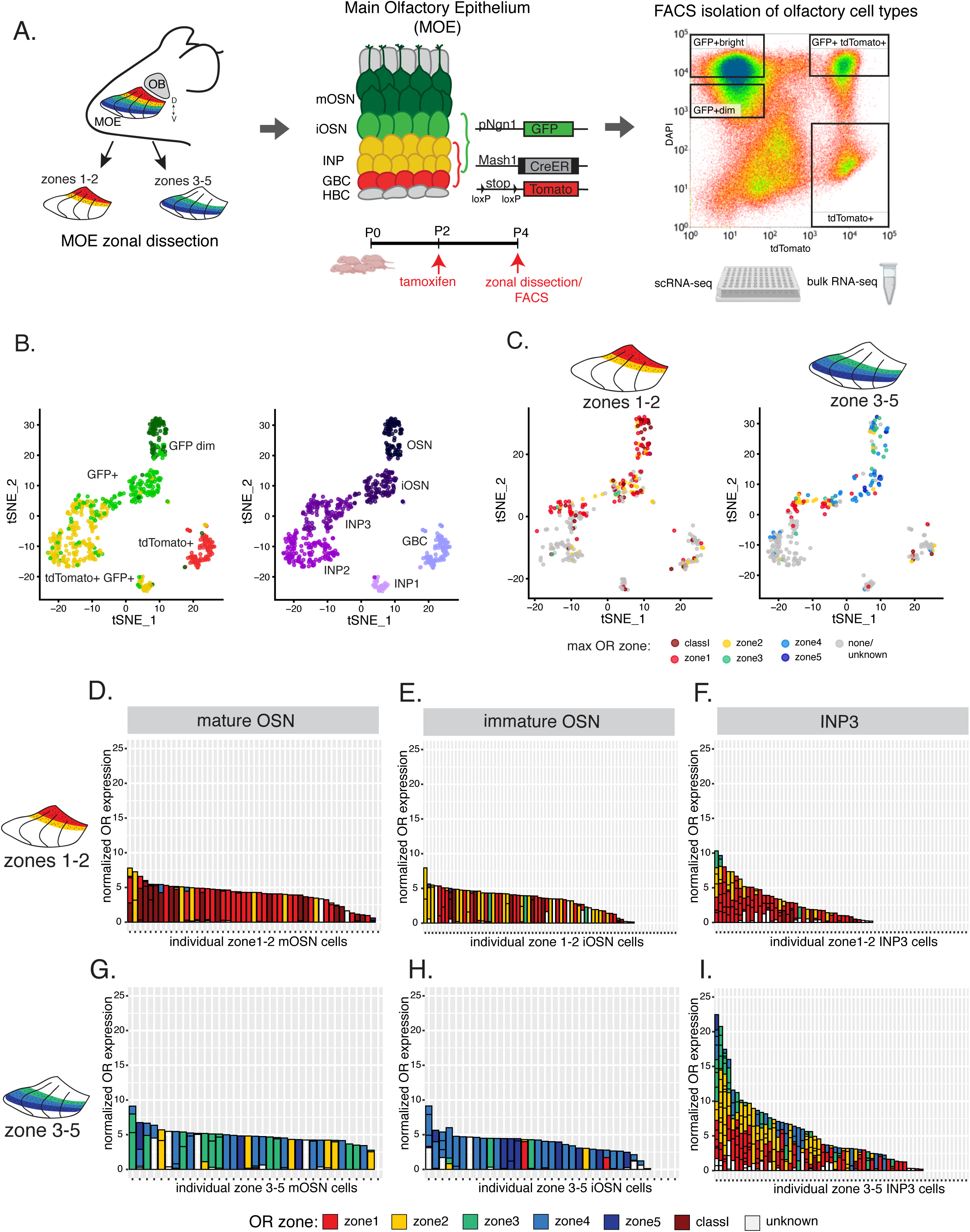
Single cell RNA-seq analysis reveals that developing olfactory progenitors undergo a timed transition in their repertoire of OR expression. (A) Schematic of the main olfactory epithelium (MOE) and the genetic and experimental strategy for isolating four cell populations at different stages of olfactory sensory neuron development (GBCs, INPs, iOSNs and mOSNs) from the same tissue. A representative FACS plot is shown. GBC, globose basal cell; INP, immediate neuronal precursor; iOSN, immature olfactory sensory neurons; mOSN, mature olfactory sensory neuron. (B) tSNE clustering of FAC-sorted cell populations with Seurat based on the most variable genes showing the separation of single cells into 6 populations. The 6 populations were assigned cell identities using expression of known MOE markers (See also Figure S1). Clustering shows the relationship between the FAC-sorted populations (left) and cell lineage (right). (C) tSNE clustering from B, separated by zonal origin colored according to most highly expressed OR. Left: cells isolated from zone1-2 dissected MOE; Right: cells isolated from zone3-5 MOE. (D) OR expression in single cells from zone 1-2 MOE (D,E,F) and zone 3-5 MOE (G,H,I). Y axis shows OR expression in normalized UMI counts for different ORs (separated by black lines) within individual cells (shown on the x axis). ORs are colored according to their zonal index (as annotated in(Tan and Xie, 2018)

To further characterize this intriguing zonal transition, we determined the zonal identity of every OR mRNA detected in each cell, using an expression threshold of 3 unique molecular identifiers (UMI) per OR. Consistent with previous reports, mOSNs predominantly express a single OR with a zonal identity that is in accordance with the zonal origin of the cell, i.e. zone 1-2 ORs in dorsal dissections and zone 3-5 ORs in ventral dissections (Figure 1D). We find that iOSNs also express a single OR with the correct zonal identity (Figure 1E). However, in INP3 cells, where polygenic OR expression is common, there is a striking departure from zonal restrictions in the ventral zones. While INP3 cells from dorsal zones co-express almost exclusively zone 1-2 ORs, INP3 cells from ventral zones co-express ORs from every zone, with zone 1-2 ORs being the most common (Figure1F, Supplemental Figure S1B). Specifically, we detect zone 1-2 ORs in 43 INP3 cells from ventral zones, and 3-5 ORs in 29 of them, with only 5 INP3 cells expressing exclusively zone 3-5 ORs. This deviation from zonal restrictions cannot be attributed to dissection errors, since both iOSNs and mOSNs from the same dissections express ORs from the correct zonal indexes. Moreover, this result cannot be explained by lateral migration of INP cells between zones, since OSN progenitors appear spatially restricted within their zone, in which they migrate vertically towards more differentiated apical layers(Chen et al., 2004; Coleman et al., 2019).

To obtain higher zonal resolution and to independently confirm the observations afforded by scRNA-seq, we trisected MOEs and performed bulk RNA-seq in INPs and mOSNs from zones 1, 2/3 and 4/5. This approach confirms that zone 1 ORs are expressed by INPs regardless of their zonal location, whereas ORs with zone 3-5 identities are only abundant in ventrally located OSNs and not in their INP progenitors (Supplemental Figure S1C). Intriguingly, even when focusing on ORs from the same zonal index, ORs most frequently expressed in INPs are less frequently chosen in OSNs, suggesting that early, low level OR transcription in mitotically active cells is not promoting OR gene choice in post-mitotic OSNs (Supplemental Figure S1D). In summary, although zonal differences are apparent at the early stages of OSN differentiation, commitment to a zonal identity does not take place at the same developmental stage across every zone; INP3 cells are already committed to their zonal identity dorsally but appear zonally multipotent in ventral zones. Therefore, determining what mechanism shuts off zone 1 ORs and activates zone 4/5 ORs in ventral OSNs, would provide mechanistic insight to zonal specification.

### Graded heterochromatin deposition silences dorsal OR indexes in ventral zones

We previously showed that OSN differentiation coincides with heterochromatin-mediated OR silencing(Magklara et al., 2011). If OR heterochromatinization is the reason for which zone 1 OR transcription stops ventral OSNs, then zone 1 OR genes should have stronger enrichment of heterochromatic marks than the other ORs in the whole MOE. Similarly, zone 2 ORs, which are turned off in iOSNs of zones 3-5, should have higher levels of heterochromatin than zone 3-5 ORs, and so forth. Indeed, visual inspection of H3K9me3 distribution in mOSNs isolated from the whole MOE reveals that in “mixed” OR clusters that contain OR genes with various zonal identities zone 1 ORs have higher levels of H3K9me3 than zone 5 ORs (Figure 2A). Aggregate analysis of all the ORs grouped by zonal identity, corroborates the gradual reduction of H3K9me3 from zone 1 to zone 5 ORs (Figure 2B). The only exception from this pattern are the dorsally expressed class I ORs. OSNs expressing the class I ORs are intermixed with zone 1 OSNs, but are a distinct subpopulation that relies on divergent gene regulatory mechanisms to control OR expression (Enomoto et al., 2019; Hirota et al., 2007).

**Figure 2:**
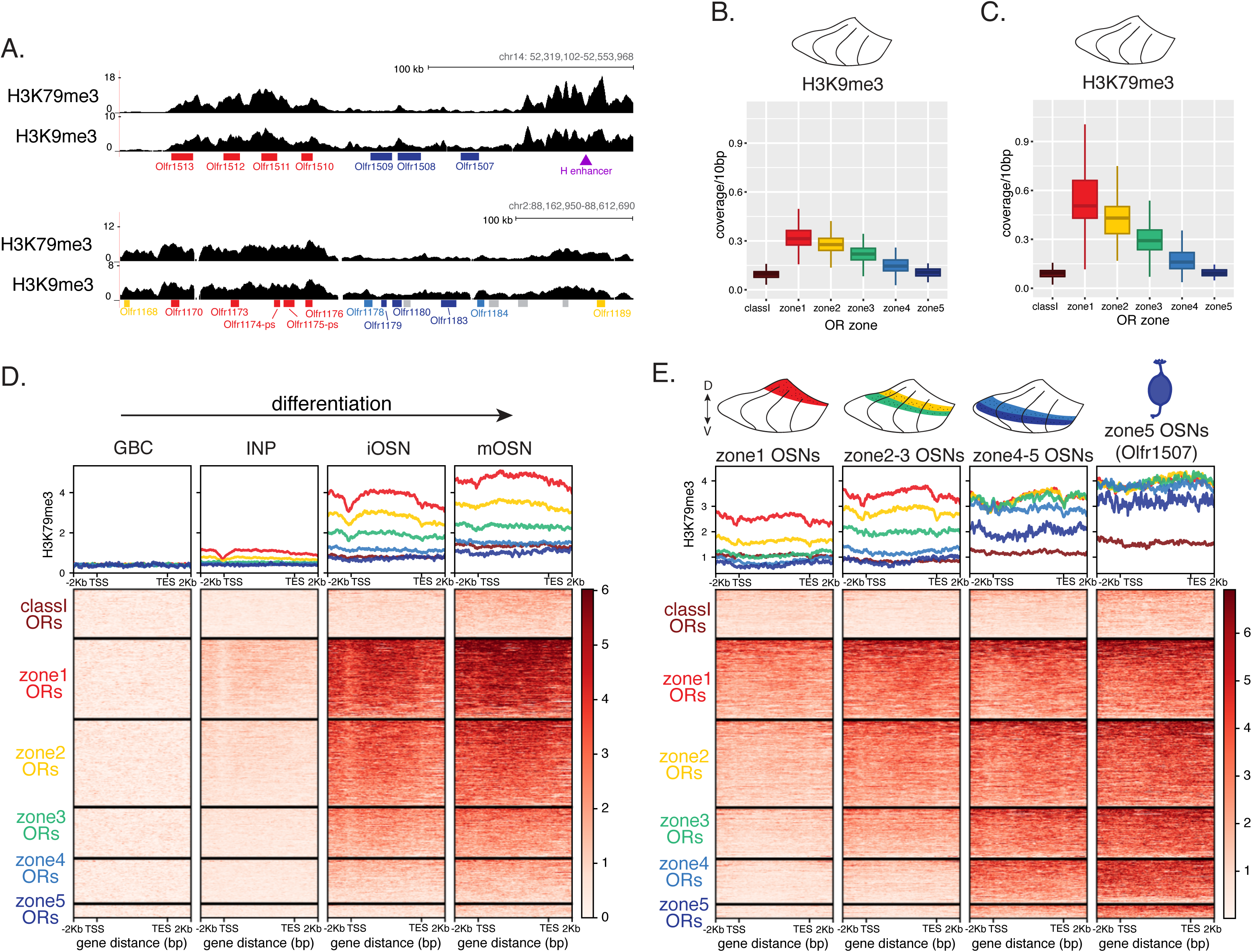
Heterochromatin deposition silences OR genes from lower zones. (A) Signal tracks of H3K9me3 and H3K79me3 native ChIP-seq from the whole MOE over two representative OR clusters. Below the signal track OR genes are colored according to their zonal index: zone1 ORs in red, zone2 ORs in yellow, zone 5 ORs in blue and ORs with unknown zone in gray. Both histone marks also flank the H Greek Island enhancer (purple triangle). (B) H3K9me3 native ChIP-seq in the whole MOE. Box plots of read density over OR gene bodies separated by their zonal index depict a pattern of deposition that is high on zone 1 OR genes, progressively decreases with higher zonal OR indexes, and is absent on class I ORs. (C) The same analysis as in B but for H3K79me3 native ChIP shows an even stronger zonal pattern of deposition. (D) H3K79me3 native ChIP seq in GBC, INP, iOSN and mOSN populations (isolated by FAC-sorting cells as described in Figure 1A) shows onset of H3K79me3 deposition in INP cells. Each row of the heatmaps shows coverage over an OR gene body (separated into categories by their zonal index). (See also Supplementary Figure S2B for H3K9me3 heatmap). (E) H3K79me3 native ChIP-seq in mOSNs from dissected zones and a pure population of Olfr1507 (zone 5 OR) expressing mOSNs. Colored MOE schematics above each heatmap depicts the zone of dissection. (See also Supplementary Figure S2C for H3K9me3 heatmap).

Searching for additional histone marks with zonal correlations, we identified an even stronger zonal influence in the distribution of H3K79me3, the enrichment levels of which decline sharply across more ventral OR identities (Figure 2A, C). H3K79me3 is an intriguing core histone modification that may play opposing roles in gene regulation(Farooq et al., 2016). It was originally described as a heterochromatic mark enriched on mammalian pericentric and subtelomeric repeats, with an active role in establishment of heterochromatin and transcriptional silencing (Jones et al., 2008). However, H3K79me3 may also play a role transcriptional activation(Steger et al., 2008), as it is enriched in developmentally regulated and cell type specific enhancers(Bonn et al., 2012; Godfrey et al., 2019). Consistent with this, we previously showed that Greek Islands are surrounded by high levels of H3K79me3, in such a specific fashion that it was used as a criterion for the distinction of OR enhancers from other OSN enhancers(Markenscoff-Papadimitriou et al., 2014). Interestingly, the levels of H3K79me3 and H3K9me3 at the sequences flanking OR enhancers are similar between OSNs from different zones but follow the general trend of gradual increase towards more ventral zones (Supplemental Figure S2A).

A second prediction from our scRNA-seq data is that deposition of heterochromatic marks should be synchronous with the transcriptional commitment to a zonal identity, i.e. should be occurring precisely at the INP to iOSN transition. To define the timing of heterochromatin formation, we performed ChIP-seq for H3K9me3/H3K79me3 on FAC-sorted GBCs, INPs, iOSNs, and mOSNs from the whole MOE. This analysis revealed that H3K9 and H3K79 trimethylation is first detected at low levels in INPs, specifically on zone 1 ORs. Since we don’t have genetic markers to sort specific INP subpopulations, we postulate that the appearance of this weak signal could coincide with the onset of OR coexpression in INP3 cells. Consistent with these marks being deposited at more differentiated stages, the levels of both H3K9me3 and H3K79me3 become stronger in iOSNs, the cell stage at which zone 1 ORs are transcriptionally replaced by ORs with the correct zonal identity (Figure 2D, Supplemental Figure S2B). Moreover, as in mOSNs, we detect intermediate enrichment of H3K9me3/H3K79me3 on zone 2 and 3 ORs and lowest levels of heterochromatic marks on zone 4/5 ORs.

The distribution of heterochromatic marks across the OR sub-genome suggests that ORs from dorsal zonal indexes may become increasingly silenced as we move to more ventral zones. To test this, we performed ChIP-seq in mOSNs isolated from micro-dissected MOE zones 1, 2/3 and 4/5. This experiment revealed the following recurrent pattern: In each zonal dissection ORs from more dorsal indexes have the highest levels of H3K9me3/H4K79me3; ORs from the correct zonal index have intermediate levels; and ORs from more ventral zonal indexes have the lowest levels of these two modifications (Figure 2E, Supplemental Figure S2C). In accordance to this trend, within tissue dissections spanning two zones we consistently observe higher levels of H3K9me3/H3K79me3 on OR genes from more dorsal indexes: zone 2 ORs have higher signal than zone 3 ORs in the zone2/3 dissection, and zone 4 ORs have higher signal than zone 5 ORs in the zone 4/5 dissection. However, these differences fade in more ventral zones, as zone 1-3 OR indexes have equally high levels of heterochromatin in zones 4/5. In fact, if we perform the same analysis in OSNs obtained exclusively from zone 5 mOSNs, which we achieved by FAC-sorting OSNs expressing the zone 5 OR Olfr1507, we find that ORs from all 5 indexes have similar levels of H3K79me3 (Figure 2E), except of the transcriptionally active Olfr1507 (Supplemental Figure S2D). Interestingly, H3K79me3 ChIP-seq in iOSNs from zones 4/5 shows that enrichment of H3K79me3 on zone 4/5 ORs is lower than on zone 1-3 ORs (Supplemental Figure S2E). Thus, although zone 4/5 ORs eventually obtain high levels of H3K79me3 within their expression zone, these enrichment levels are reached after the completion of OR choice and only at the ORs that were not chosen for transcription. In contrast, ORs from zone 1-3 already have high H3K79me3 enrichment levels from the iOSN stage in the ventral-most zones and are likely excluded from the process of OR gene choice. This pattern explains why ventral OSNs do not express ORs with dorsal indexes, but it raises the reverse question: What prevents dorsal OSNs from choosing ventral ORs that have low levels of repressive chromatin marks?

### Graded genomic compartmentalization precludes expression of ventral ORs in dorsal zones

OR gene activation requires formation of OR compartments, assembly of a multi-enhancer hub next to the OR compartment, and association of this hub with one of the proximal OR alleles. To test if zonal positioning influences this process, we performed *in situ* Hi-C in FAC-sorted OSNs from zones 1, and 4/5, as well as OSNs expressing OR Olfr17, representing zone 2 OSNs. This analysis revealed that long-range *cis* and *trans* genomic interactions between ORs are strongly linked to the zonal index of each OR and the zonal identity of the OSN, mimicking the deposition of heterochromatin. For example, inspection of the genomic interactions between 3 OR clusters in chromosomes 2, shows that a cluster that contains mostly zone 4/5 ORs makes contacts with the other two clusters only in ventral OSNs. In contrast, the other 2 OR clusters, which are either enriched for zone 1-3 ORs, or contain ORs from every zonal index, make strong genomic contacts with each other in all three zonal OSN populations (Figure 3A). To expand this analysis to every OR, we measured the frequency of interchromosomal OR-OR interactions across zone 1, 2-3, and 4-5 OR indexes in the 3 zonal OSN populations, reaching a similar conclusion: In each zonal OSN identity ORs from more dorsal indexes have the highest levels of *trans* Hi-C contact; ORs from the correct zonal index have intermediate levels; and ORs from more ventral zonal indexes have the lowest levels of genomic contacts (Figure 3B). Intriguingly, we observe a consistent Hi-C contact frequency for the set of OR genes competent to be chosen across different anatomical zones (Figure 3B). We detect a similar pattern in iOSNs, with differences in genomic contacts between different OR identities being sharper than in mOSNs, especially in *trans* (Supplemental Figure S3A, B). At this developmental stage there is a strong correlation between the levels of H3K79me3 and the frequency of Hi-C contacts (Supplemental Figure S3C), suggesting that the two processes may be mechanistically linked, as recently proposed for heterochromatin and genomic compartmentalization(Falk et al., 2019; Mirny et al., 2019).

**Figure 3:**
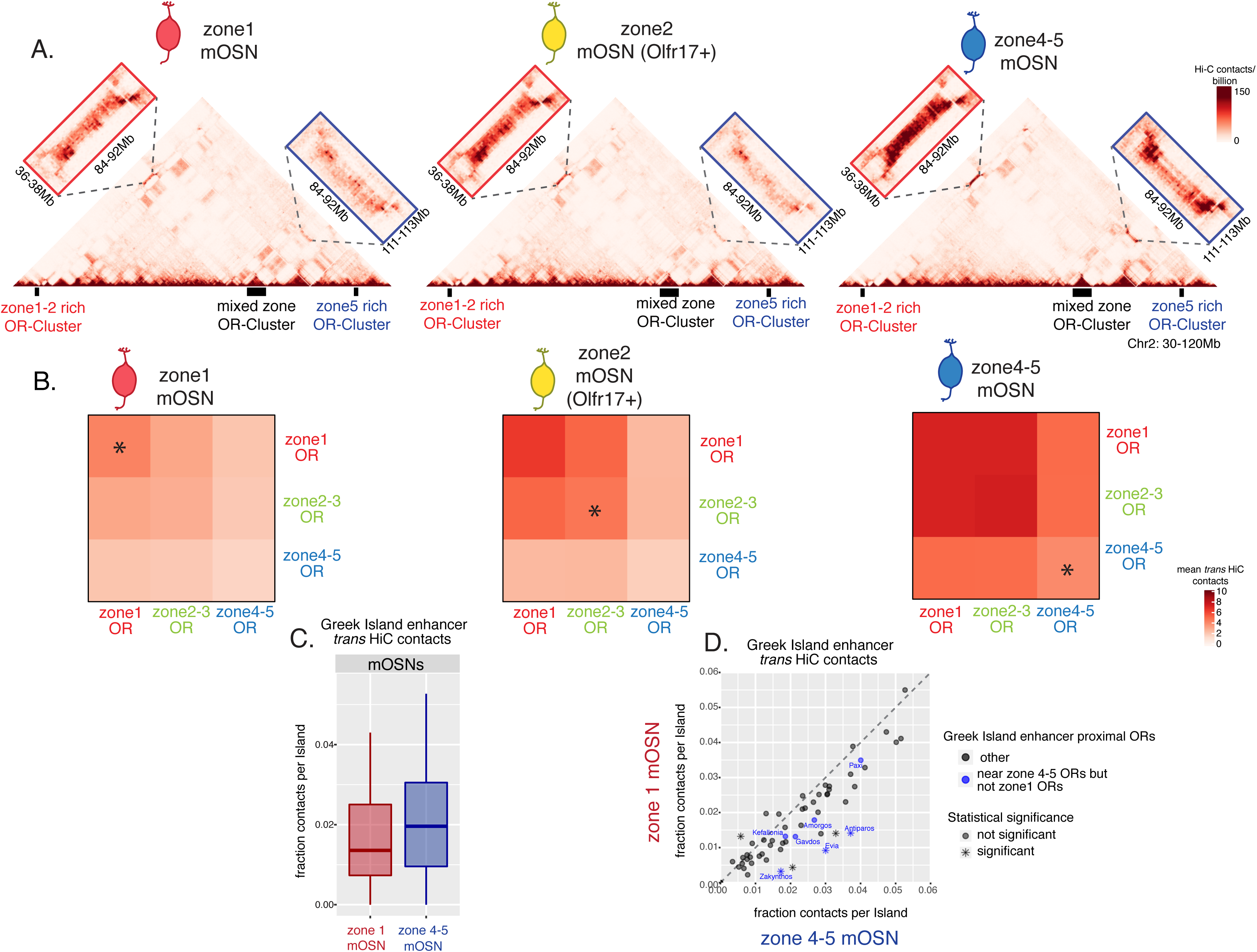
Zonal OR compartmentalization permits OR genes from more ventral zones to be recruited into the OR compartment. (A) In situ Hi-C contact matrices of a 90Mb region of chromosome 2 from zone1, zone2 and zone 4/5 mOSN show long range cis interactions between 3 large OR gene clusters. The cluster on the right (blue) is enriched for zone 4/5 OR genes, does not contain any zone1 OR genes, and only makes strong long-range interactions with the other clusters in zone 4-5 mOSNs. Zone1 and zone4/5 mOSNs were isolated from zonally dissected olfactory epithelium and zone2 mOSNs were isolated from Olfr17 (a zone2 OR) reporter mice, published in(Monahan et al., 2019). (B) Heatmaps of average interchromosomal Hi-C contacts between OR genes annotated by their zonal index at 50Kb resolution (excluding bins that contain ORs genes expressed in multiple zones and Greek Island enhancers) shows increased contacts in cells from more ventral zones (zone1 mOSNs < zone2 mOSNs < zone4/5 mOSNs). OR genes have a similar frequency of contacts in the mOSN population where they are expressed (zone1 ORs in zone1 mOSNs, zone2-3 ORs in zone 2 mOSNs and zone4-5 ORs in zone4/5 mOSNs), marked with an asterisk. (C) Quantification of Greek Island enhancer interchromosomal interactions show Greek Islands make a similar fraction of their total H-C contacts with other Greek Islands in zone1 and zone4/5 mOSNs. (D) Analysis of contacts made by individual Greek Islands show that 6 out of 59 Greek Islands have differential frequency of interchromosomal interactions between the zone1 and zone5 mOSNs: 5 form significantly more interactions in zone 4/5 cells and one in zone1 cells (Wilcoxon rank sum test: p-value < 0.05). Greek Islands residing in OR clusters with zone 4-5 ORs and no zone1 ORs (blue) appear relatively less engaged in zone1 mOSNs. All Hi-C analysis was performed at 50kb resolution and shows merged data from two biological replicates that yielded similar results when analyzed separately.

We then asked if there are zonal differences between the long-range contacts made by Greek Islands. Consistent with the small variations in flanking H3K79me3/H3K9me3, t*rans* Hi-C contacts between Greek Islands are comparable between zones 1 and 4/5 mOSNs, with a small increase towards zone 4/5 (Figure 3C). This is consistent with results from FAC-sorted OSNs expressing Olfr16 (zone1 OR), Olfr17 (zone2 OR), and Olfr1507 (zone5 OR), showing similar patterns of active OR-Greek Island interactions between the 3 zones(Monahan et al., 2019). One exception is detected on enhancers residing in OR clusters that contain predominantly zone 4-5 ORs. As these OR clusters make very infrequent contacts with other OR clusters in zone 1 (Figure 3A), enhancers located in these clusters also have reduced contacts with distant Greek Islands, consistent with the notion that OR compartmentalization promotes enhancer compartmentalization (Supplemental Figure S3D, E).

To obtain insight to the zonal differences in OR compartmentalization at single cell resolution, we performed Dip-C, a high coverage version of single cell Hi-C, in 96 mature OSNs, 48 each from zones 1 and 4/5. To retain allelic information, we crossed OmpiresGFP mice (mixture of Bl6/123 strains) to Castaneous (Cas) mice and used Cas-specific SNPs to distinguish Cas from non-Cas alleles. After filtering for high quality cells, we excluded 4 cells and generated contact maps with a median of over 400,000 contacts per cell. We used the haplotype resolved data to compute distances of all genomic loci at 20kb resolution and generated 3D models for all cells, as previously described(Tan et al., 2019). In these models, we were able to detect formation of interchromosomal compartments in mOSNs from both zones (Figure 4A, B). Analyzing distances between pairs of OR loci in the 3D model we determined the size and complexity of OR compartments in each cell. Consistent with predictions from our bulk Hi-C, ventral 4/5 OR compartments are larger and contain more ORs from more chromosomes in zone 4/5 OSNs than zone 1 OSNs (Figure 4C, D, Supplemental Figure S4A-D). Despite the fact that there are zone-specific features of OR compartmentalization, within a zonal mOSN population every cell has a distinct pattern of long range *cis* and *trans* OR contacts (Supplemental Figure S4A, B). As previously shown, Greek Islands have stronger interchromosomal contacts than OR genes, that are less influenced by zonal identity than OR-OR interactions (Supplemental Figure S4E, F). Thus, in a given zone4/5 OSN, on average around 27% of enhancers were engaged in interchromosomal contacts within 2.5 particle radii compared to 22% in zone1. But the size of the largest hub (ranging from 4-8 enhancers), presumably the site of OR transcription, was comparable in the zone1 and zone4/5 cells (Supplemental Figure S4G). In general, Greek Islands make strong contacts with ORs from every zone, however in zone 4/5 mOSNs they have a significant preference for zone 4-5 ORs compared to zone 1-3 ORs. We do not detect a significant association of Greek Islands with zone 1 ORs in zone 1 mOSNs, most likely because ∼50% of these cells express class I ORs, which are not regulated by the multi-enhancer hub.

**Figure 4:**
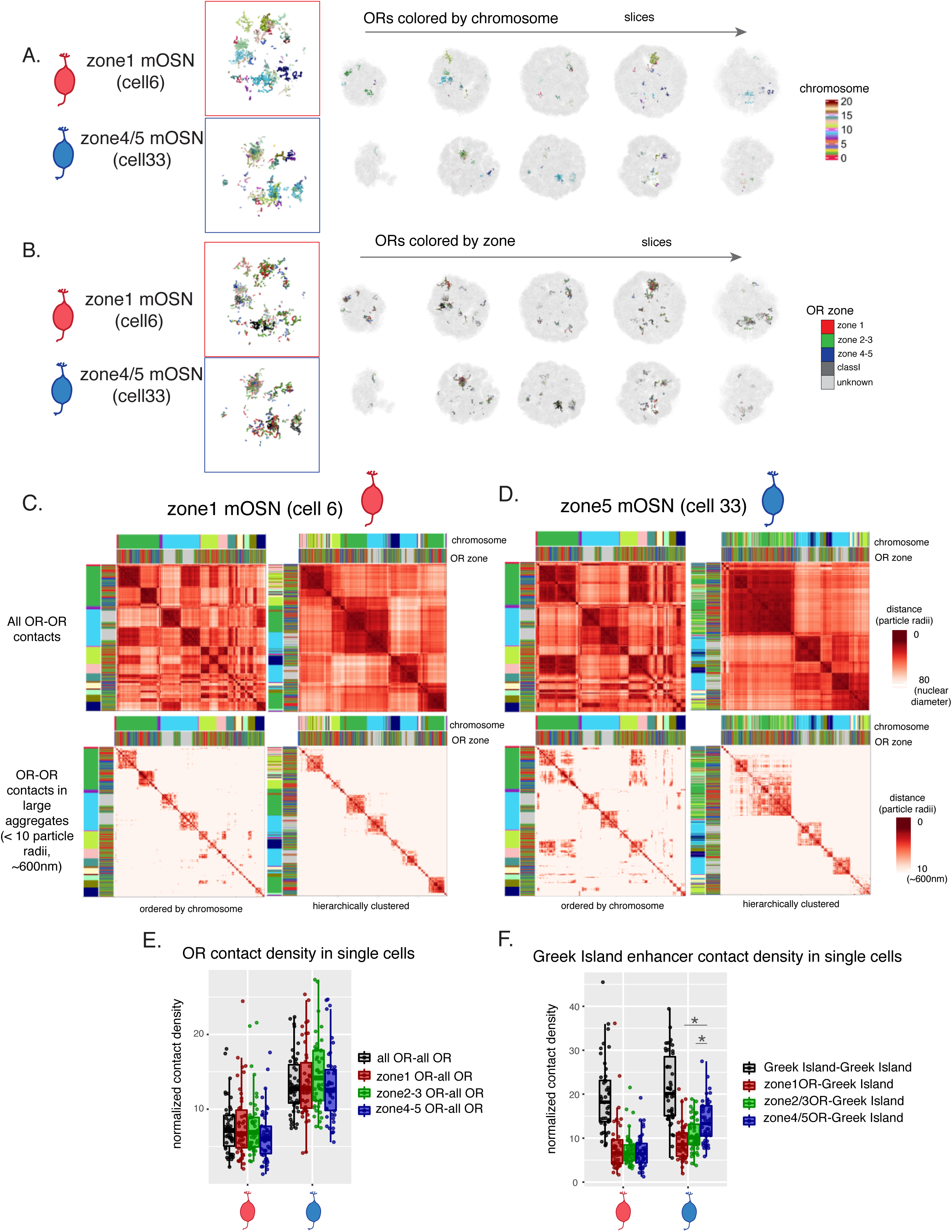
Diploid chromatin conformation capture (Dip-C) reveals differences in OR compartments in single zonal mOSNs. (A, B) 3D genome structures in single cells generated from Diploid chromatin conformation capture (Dip-C) in 96 mOSNs--48 cells from each zone1 and zone4/5. Cells were isolated from OMP-ires-GFP mice crossed to Castaneous. (A) Representative 3D structures of OR compartments in a zone1 and a zone4/5 mOSN colored by the chromosome, shows increased complexity of OR compartments in the zone 4/5 mOSN. Serial sections at 15 particle radii (approximately ∼900nm) show several OR aggregates in each cell. (B) The same 3D structures in (A) colored by zonal OR annotation of each particle (analogous to 20kb of chromatin) shows OR genes of all zonal indexes intermingling. (C, D) Single cell heatmaps of pairwise distances between OR genes generated from 3D nuclear structure in a zone1 mOSN (C) and zone4/5 mOSN (D) show an intermingling of OR genes from different chromosomes. Heatmaps sorted by chromosome order (right) or via hierarchical clustering (left) show increased interchromosomal contacts in the zone4/5 mOSN. The heatmap displaying contacts at any distance (top) and contacts within 10 particle radii (approximately ∼600nm) shows larger aggregates in the zone 4-5 mOSN. Heatmaps of contacts within 10 particle radii are shown for all cells in Supplementary Figure S4. Only OR genes for which the zonal annotation is known (and that do not reside within 20kb of an OR of another zonal index or within 50kb of a Greek Island) were used to generate the heatmaps. (E) Analysis of contact densities of interchromosomal contacts of all ORs (black), zone1 ORs (red), zone 2/3 ORs (green) and zone4/5 ORs (blue) with other ORs confirms that zone4/5 mOSNs (blue cell) have an increased density of interchromosomal contacts between OR genes of all zonal indexes, relative to zone1 mOSNs (red cell). (F) A similar comparison of Greek Island contact densities with all ORs (black), zone1 ORs (red), zone2/3ORs (green) and zone4/5 ORs (blue) shows that in zone4/5 cells Greek Islands form stronger interchromosomal interactions with zone4-5 OR genes relative to zone1 and zone2-3 OR genes (Wilcoxon rank sum test: p-values = 1.078×10-6 and 0.002, respectively).

### Graded expression of NFI paralogues implicates them in zonal OR gene regulation

The graded increase in the deposition of H3K9me3/H3K79me3 and OR compartmentalization from dorsal to ventral zones has important regulatory implications. Instead of deploying distinct, zone-specific transcription factor combinations, developing OSNs could establish zonal restrictions with shared factors expressed in a graded fashion. The transcriptional regulators with an established role in OR gene choice are expressed at similar levels in cells isolated from different zones, with the only exception being a somewhat lower expression of Ldb1 in zone 1 (Supplemental Figure 5A). Thus, we searched for additional transcription factors that may specifically contribute to the zonal restrictions on OR gene choice. Taking into account the developmental onset by which zonal differences appear, we searched for transcription factors that have strong expression in both INPs and OSNs that is graded across the dorsoventral axis of the MOE. We identified numerous transcription factors that are expressed in a zonal fashion at various stages of OSN differentiation, some of which were previously described (Duggan et al., 2008; McIntyre et al., 2008; Parrilla et al., 2016; Tietjen et al., 2005), and some, like the zone 1 enriched Nhlh2, Hey1 and Nr2f2, being associated with zonal expression for the first time(Figure 5A-D and Table 1). Importantly, from all the transcription factors with 3fold expression difference between zones 1 and 4/5, the NFI paralogues NFIA, B, and X are the only ones that retain strong, zone-influenced expression from the early INP to the mOSN stage (Figure 5A-D). These three members of the nuclear factor I (NFI) family of transcription factors control a plethora of developmental and cell specification processes(Gronostajski, 2000; Zenker et al., 2019), and were previously implicated OSN differentiation(Baumeister et al., 1999; Behrens et al., 2000). NFIs are expressed at higher levels in zone 4/5 cells, from INP to mOSN, with NFIA and NFIB being expressed higher in INPs and iOSNs, and NFIX being upregulated in mOSNs (Figure 5E, Supplemental Figure 5B).

**Figure 5:**
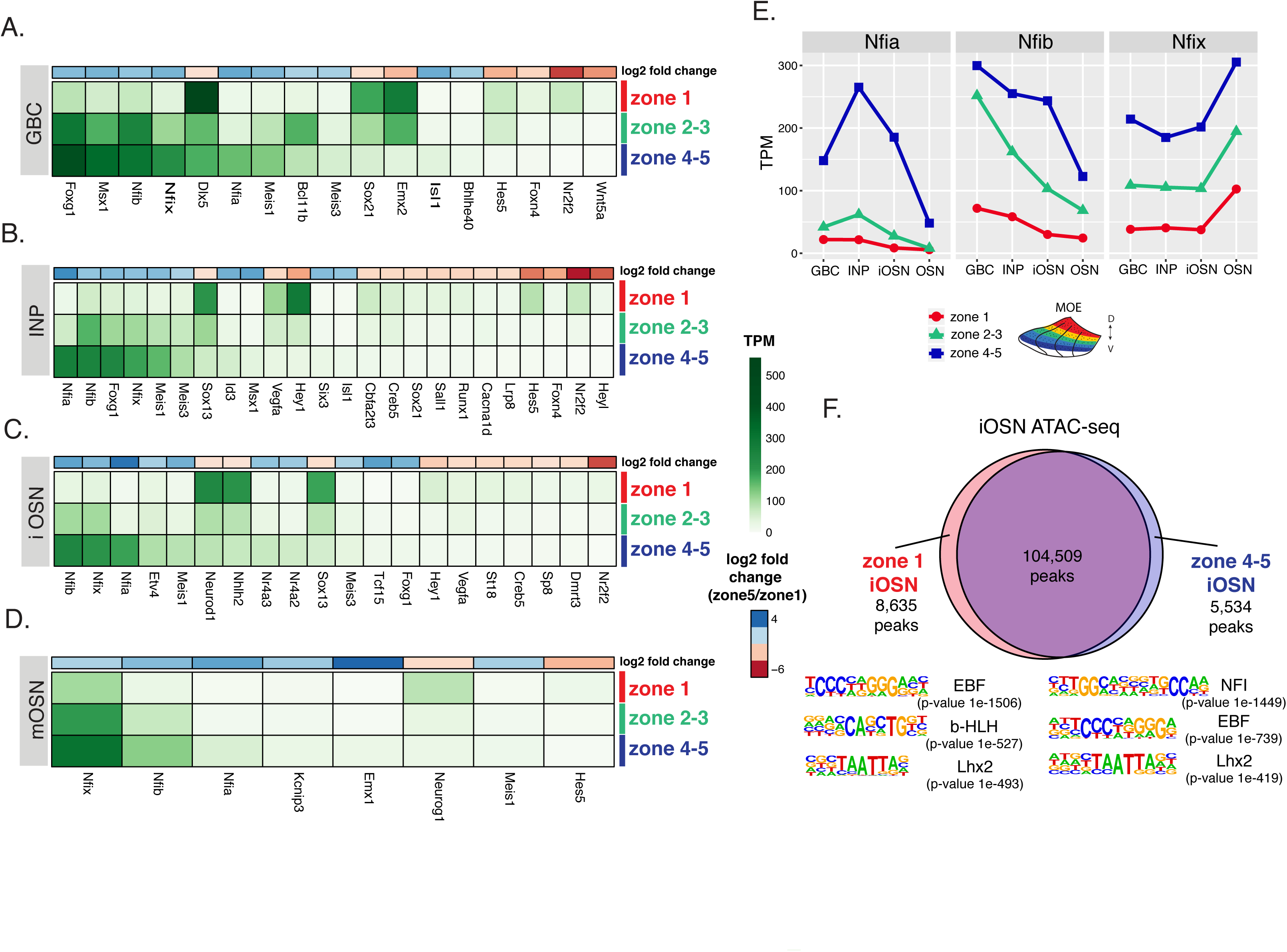
NFI family transcription factors have a graded expression in cells from different zones. (A, B, C, D) Heatmaps showing the expression levels of differentially expressed transcription factors between zone1, zone 2/3 and zone 4/5 cells at different stages of development: GBC, INP, iOSN and mOSN. The shown transcription factors are significantly differentially expressed between zone5 and zone1 cells with an adjusted p-value of <0.05, at least a three-fold change in expression, and an expression of at least 15 TPM. A broader list of transcription zonal factors is included in Table1. The heatmaps is sorted based on expression in zone 4/5 cells and a color bar above each heatmap shows the log2 fold change in zone 4/5 cells relative to zone 1 cells (blue values represent genes higher in zone 4/5). (E) Expression levels are shown for each of the members of the NFI family: NFIA, NFIB, and NFIX (NFIC, which is not expressed in the MOE, is excluded). Plots of expression at the four developmental stages in zone 1 cells (red), zone 2/3 cells (green) and zone 4/5 cells (blue) show higher levels of transcription of all three factors in zone 4/5 and show different patterns of expression during the course of development (NFIA and NFIB decrease in mOSNs, while NFIX increases). (F) Pie chart showing the overlap of Omni-ATAC peaks in zone1 and zone 4/5 iONS and the top de novo motifs identified with HOMER in differentially accessible peaks (p-value < 0.05). The NFI motif is the most highly enriched motif in zone4/5 iOSN unique peaks.

Although the continuous zone 4/5 enrichment of NFI genes prioritizes their functional interrogation over other zonally expressed transcription factors, we sought additional evidence for a functional role in the specification of ventral zones. First, we performed omni-ATAC(Corces et al., 2017) on iOSNs from zones 1 and 4/5. This experiment revealed the existence of >8,000 zone1-, and >5,000 zone 4/5-specific peaks. Lhx2 and Ebf motifs are abundant in both peak groups (Figure 5C), consistent with the key regulatory function of these two transcription factor types in OSN identity. Moreover, zone1-specific peaks have enrichment for bHLH motifs in agreement with the increased expression of numerous bHLH transcription factors in zone 1 iOSNs (Figure 5A, C). Interestingly, the most significantly enriched zone-specific motif overall is NFI, which is identified in the zone 4/5 peaks, further supporting a functional role of zonal NFI expression (Figure 5C). If we focus our omni-ATAC analysis on OR promoters, which have weak ATAC signal that for most promoters does not reach peak-calling threshold, we detect stronger signal on zone 1 promoters in both zone 1 and 4/5 iOSNs. Interestingly, as is the case for H3K9me3/H3K79me3 distribution and OR compartmentalization, ATAC signal is increased on most OR promoters in zone 4/5 iOSNs, and even becomes apparent on zone 4/5 promoters, although these remain the weakest group. Although there is no enrichment of bHLH motifs on zone 1 OR promoters compared to zone 4/5 promoters, NFI motifs are twice as frequent on zone 4/5 promoters (Supplemental Figure S5C). It should be noted zone 5 promoters have low quality Ebf motifs, suggesting the possibility that NFI binding compensates for weaker Ebf binding on these sequences (Supplemental Figure S5D).

To further investigate the genome wide binding of Nfi proteins we performed CUT&RUN(Meers et al., 2019a; Meers et al., 2019b) with an antibody against Nfia protein. We obtained *∼*2,000 peaks genome wide, with NFI motifs significantly enriched in these peaks, but we did not detect Nfi binding on zone 4/5 OR promoters (data not shown). This, however, is not surprising, since due to cellular heterogeneity none of the transcription factors known to regulate OR expression can be detected on OR promoters (Monahan et al., 2017). However, we detect Nfia signal on Greek Islands despite the lack of full NFI motifs on these enhancers (Supplemental Figure S5E). Consistent with binding of additional factors, like Nfi proteins, to the core of Lhx2/Ebf/Ldb1, Greek Islands in zone 4/5 iOSNs have increased ATAC-seq signal, especially residing in OR clusters containing predominantly zone 4/5 ORs (Supplemental Figure S5F-G). These observations, taken together, suggest that Nfi proteins may contribute to the aforementioned zonal differences of heterochromatin assembly, OR compartmentalization, and Greek Island hub formation, and they may bias association of the Greek Island hub with zone 4/5 ORs in ventral zones of the MOE. Thus, we sought a functional characterization of the role of NFIA, B, and X in zone 4/5 specification.

### NFI A, B, and X gradients regulate zonal OR expression

To interrogate the potential role of NFI A, B and X in zonal OR expression, we adopted a loss-of-function approach. To overcome possible redundancy between the NFI paralogues, we deleted all three genes simultaneously. To assure complete removal of Nfi activity before the INP-to-iOSN transition we used the Krt5CreER driver, which is expressed in HBCs, the quiescent stem cells of the MOE. We crossed Krt5CreER; tdTomato mice to NFIA,B,X fl/fl mice(Clark et al., 2019), and induced recombination with tamoxifen. To force the quiescent HBCs to differentiate into OSNs, we ablated the MOE with methimazole and allowed 40 days for a complete restoration by the marked progeny of the NFI KO or control HBCs (Supplemental Figure S6A), as previously described(Monahan et al., 2019). We first performed IF in MOE sections of control and triple NFI KO mice using antibodies for ORs Olfr1507 (zone 5 OR) and Olfr17 (zone 2 OR). This analysis revealed an almost complete depletion of Olfr1507-expressing OSNs (Figure 6A), and reciprocal ectopic expression of Olfr17 in zone 5 OSNs (Figure 6B). To analyze expression of all OR genes, we FAC-sorted tdTomato^+^ control and KO OSNs from the whole MOE and performed RNA-seq. Consistent with the immunofluorescence imaging results, we detect a strong reduction of all zone 4/5 OR mRNAs and a concomitant increase of zone 2/3 mRNAs in the triple KO OSNs (Figure 6C). To explore how stable zonal OR specification is after it is established, we deleted the 3 NFIs with OMPiresCre, a driver that induces recombination in mature OSNs (Supplemental Figure S6A). Intriguingly, triple NFI deletion in mOSNs has no effects on zonal OR expression, with most zone 2/3 and 4/5 OR mRNAs being as abundant as in control mOSNs (Figure 6C). In contrast to ORs, numerous non-OR zone 4/5 markers are downregulated in both the “early” and “late” NFI deletion (Supplemental Figure S6B). Thus, unlike other zonally expressed genes, OR patterning persists for weeks, suggesting that heterochromatinization and genomic compartmentalization constitute forms of long-term nuclear memory in post-mitotic mOSNs.

**Figure 6:**
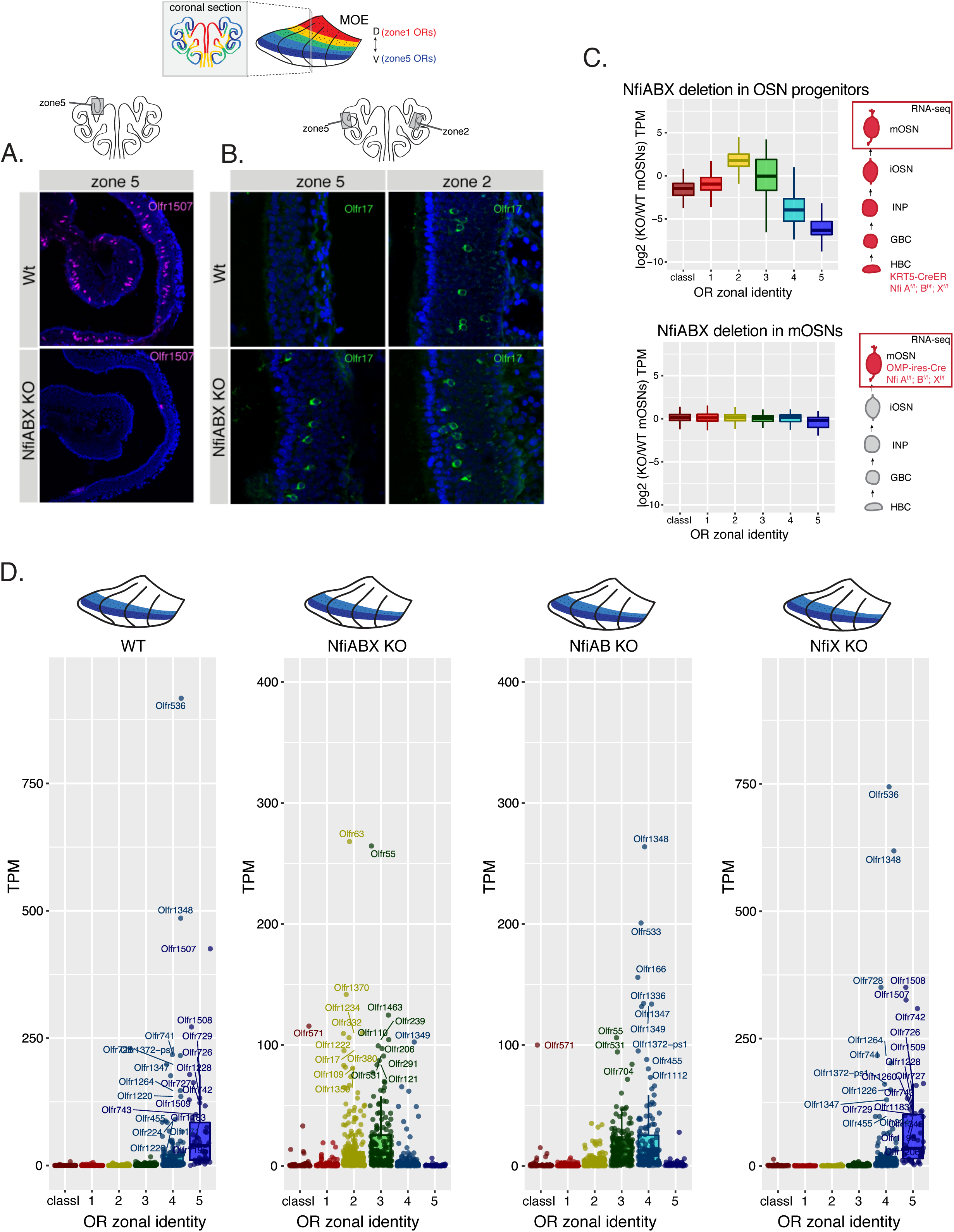
Triple knockout of NFI ABX in progenitors transforms zone4/5 olfactory epithelium into zone2/3. (A) Olfr1507 (a zone5 OR) immunofluorescence staining in MOE sections (magenta) of adult Nfi ABX triple knockout mice and age matched wt mice. Nuclei are stained with Dapi (blue). Images were taken in the same location, indicated on the schematic of a coronal section of the MOE. (B) Olfr17 (a zone2 OR) immunofluorescence staining in MOE sections (green) of adult NFI ABX triple knockout mice and age matched wt mice shows normal Olfr17 expression continues in NFI ABX knockout mice in zone 2, while also spreads into more ventral zones and is detected in zone 5. Images were taken in the same spot in the MOE, indicated in the schematic. Nuclei are stained with Dapi (blue). (C) Comparison of OR gene expression in NFI ABX triple knockout and wt cells from the whole MOE. NFIs are deleted either in progenitors (top) using a KRT5-CreER driver or in mOSNs (bottom) using an OMP-ires-Cre driver (as described in Supplementary Figure S6A). (D) OR expression in NFI ABX triple knockout, NFI AB double knockout, NFIX knockout and wt mOSNs from zone4/5 dissected MOE. Knockout was induced in progenitors (according to the “early” knockout strategy described Supplementary Figure S6A). Plots show a different pattern of transcription of OR genes in the different genotypes. Quantification of differentially expressed ORs in each genotype is shown in Supplementary Figure S6D.

The aforementioned observations suggest that zone 2/3 ORs substitute zone 4/5 ORs in the ventral-most zones of the “early” triple NFI KO. To directly test this prediction, we performed RNA-seq analysis in OSNs isolated from micro-dissected zones 4/5. This experiment confirms the strong transcriptional downregulation of most zone 4/5 ORs, and the ectopic expression of many zone 2/3 ORs in the ventral-most zones (Figure 6D). This striking “dorsalization” of zone 4/5 OSNs satisfies the original criteria of homeosis (Bateson, 1894), since the overall mOSN identity is not altered by the triple NFI deletion. Only 13 out of ∼200 OSN-specific genes are significantly different between control and KO OSNs, and 117/207 non OR zone 4/5 enriched genes are still expressed in the ventral-most zones, acting as independent fiducial markers for our zonal dissection (Supplemental Figure S6C).

Having established a robust phenotype from the early triple NFI KO, we explored the redundancy between the three Nfi proteins. We analyzed separately double (NFIA, B) and single (NFIX) “early” KO OSNs. In double (NFIA, B) KO mice, zone 4/5 OSNs express predominantly zone 3/4 ORs, revealing a partial dorsal shift compared to the triple NFI KO mOSNs, which express ectopically zone 2/3 ORs (Figure 6D, Supplemental Figure S6D). A more detailed investigation revealed that while zone3 ORs are ectopically expressed in both the double and triple KO a different set of zone3 ORs are miss-expressed in each case (Supplemental Figure S6E). Thus, even within a defined zonal index, there are likely additional sub-divisions across the dorsoventral axis, influencing the relative sensitivity to the total number of deleted NFI paralogues. In this vein, zone 5 ORs with the ventral-most OR index, disappear in both the double and triple NFI KO, whereas the dorsal-most zone 1 ORs are not affected by either KO. Finally, NFIX deletion alone does not have significant consequences in the zonal expression of ORs (Figure 6D, Supplemental Figure S6D). The most likely explanation for the lack of a detectable phenotype in the NFIX KO is the fact that the other two NFI paralogues, which are more highly expressed during the critical INP-to-iOSN transition, suffice for proper zonal specification; however, as the differences between the double and triple KO indicate, NFIX has a critical contribution to this process. Notably, for each KO, the severity of the zonal transformation in OR expression correlates with the overall change in expression of non OR zone4/5 enriched genes (Supplemental Figure S6C).

### NFI A, B, and X govern heterochromatin deposition and OR compartmentalization

If the stable establishment of zonal OR patterning is mediated by heterochromatin assembly and genomic compartmentalization, then “early” triple NFI deletion should influence one or both of these processes. Deploying the genetic strategy described previously for “early” NFI deletion (Supplemental Figure S6A), we isolated control and triple NFI KO Tomato^+^ OSNs from the ventral-most zones and performed ChIP-seq and *in situ* Hi-C. ChIP-seq revealed a remarkable transformation in the pattern of H3K79me3/H3K9me3 across zonal OR identities with strong reduction of both histone modifications in zone 3-5 ORs, and little change in zone 1-2 ORs, as a whole, (Figure 7A, B), mimicking the pattern observed in zones 2/3 (Supplemental Figure S7A). Furthermore, if we group zone 2 and 3 ORs in accordance to the significance of their upregulation in zones 4/5 (Figure 7C), we make an intriguing observation in regards of H3K79me3 enrichment: zone 2 ORs that are most significantly upregulated have the lowest levels of H3K79me3 between zone 2 ORs, whereas for zone 3 ORs the trend is the opposite (Figure 7D). Furthermore, if we focus our analysis to the sub-indexes of zone 3 ORs that are ectopically expressed only in the double or only in the triple NFI KO, we observe that the former has less H3K79me3 than the latter in the triple NFI KO (Supplemental Figure S7B). These results further support the emerging concept that there is a non-linear correlation between levels of heterochromatin and transcriptional outcomes, with intermediate enrichment levels coinciding with OR gene choice.

**Figure 7:**
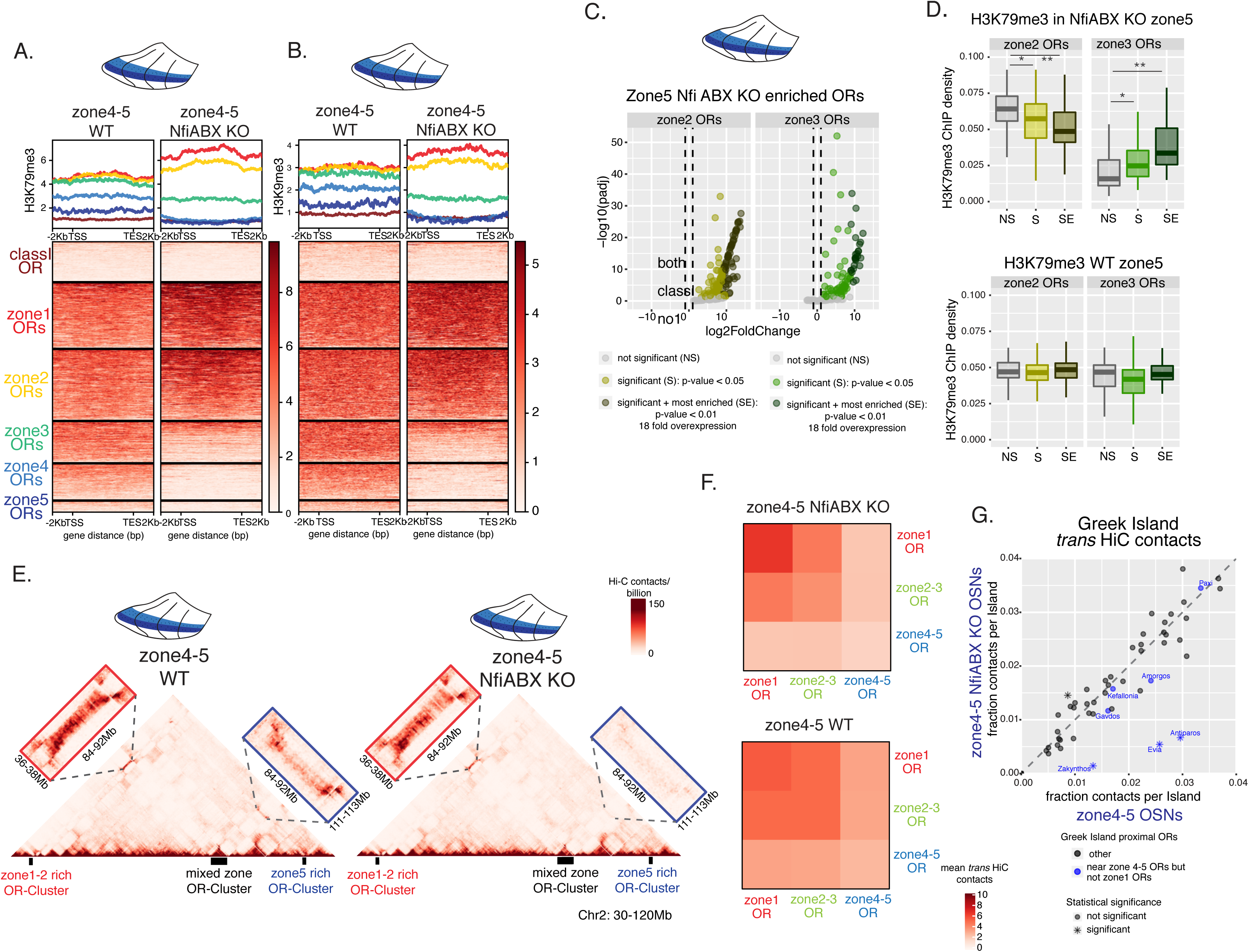
Nfi A, B and X regulate chromatin state and OR compartment formation. (A, B) Native ChIP-seq for H3K79me3 (A) and H3K9me3 (B) in Nfi ABX knockout zone 4/5 cells. Heatmaps of ChIP signal over OR gene bodies show a dramatic decrease of both histone marks on zone 3-5 ORs in NfiABX knockout compared to wt. (C) Identification of zone 2 and 3 OR genes that are ectopically expressed in RNA-seq of zone 4/5 Nfi ABX knockout olfactory epithelium. All zone 2 and zone 3 ORs were classified based on their enrichment in zone4/5 olfactory epithelium as either not significant (NS), significant (S) with an adjusted p-value<0.05, or significant & most enriched (SE) with an adjusted p-value<0.01 and a fold increase > 18. (D) Boxplots of H3K79me3 ChIP coverage over zone2-3 OR genes in Nfi ABX triple KO zone4/5 classified by their change in gene expression (top). H3K79me3 ChIP-seq data for the same genes in WT tissue (bottom). In triple KO cells there is less H3K79m3 on zone 2 SE OR genes compared to NS and S (Wilcoxon rank sum test p-values = 4.823e-07 and 0.001 respectively), and more on zone 3 SE OR genes compared to NS and S (Wilcoxon rank sum test p-values =1.649e-05 and 0.005 respectively) suggesting an intermediate level of H3K79me3 is consistent with high transcription. (E) In situ Hi-C contact matrices of a 90Mb region of chromosome 2 from wt and NFI ABX triple knockout zone 4/5 cells show long-range *cis* interactions between 3 large OR gene clusters. The cluster on the right (blue)--enriched for zone 4-5 OR genes--shows almost no long-range contacts with the other two clusters in Nfi ABX knockout Hi-C. (F) Heatmaps of average interchromosomal Hi-C contacts between OR genes annotated by their zonal index show that in Nfi ABX zone 4/5 mOSNs (bottom) zone 4/5 ORs make proportionately fewer interchromosomal contacts compared to zone 1 and zone 2/3 ORs. In contrast, wt zone 4/5 mOSNs have a more homogenous pattern of interchromosomal contacts between ORs. (G) Comparison of interchromosomal interactions made by individual Greek Islands shows that 3 Greek Islands made significantly fewer contacts in zone 4/5 Nfi ABX knockout mOSNs compared to wt (Wilcoxon rank sum test: p-value < 0.5) and one Greek Island (located in an OR cluster with one zone 3 OR) showed more contacts.

If levels of H3K79me3/H3K9me3 enrichment influence OR compartmentalization (or vice versa), then we would expect strong changes in long-range genomic interactions upon NFI deletion. Indeed, *in situ* Hi-C in control and triple Nfi KO Tomato^+^ OSNs from zone 4/5 revealed a strong reduction in the long-range genomic contacts made by zone 4/5 ORs. This reduction applies to both long range *cis* contacts between OR gene clusters (Figure 7E), as well as genome-wide *trans* interactions (Figure 7F). In fact, direct comparison of the patterns of zonal interchromosomal interactions between OSNs from the three dissected zones suggests that triple Nfi KO OSNs from zone 4/5 adopt a nuclear architecture that is most similar with the nuclear architecture of zone 2/3 OSNs (Supplemental Figure S7C). In support of this, if we plot interchromosomal OR interactions across a large mixed cluster on chromosome 2, we find that the distribution of *trans* OR contacts between ORs from different zones is quite different between zones 2/3 and 4/5; yet in the triple Nfi KO, we find that this distribution is transformed in 4/5 OSNs to an almost identical pattern with zone 2/3 OSNs (Supplemental Figure S7D). Finally, we asked if triple NFI deletion affects the frequency of *trans* contacts between Greek Islands. Overall, *trans* contacts between Greek Islands are not reduced in triple KO (Supplemental Figure S7E), consistent with the fact that we did not observe strong zonal differences for these genomic contacts. However, Greek Islands that reside on OR clusters that contain mostly zone 4/5 ORs have reduced *trans* contacts with other OR enhancers, consistent with the effects of NFI deletion on the genomic interactions of zone 4/5 ORs. Nfia CUT&RUN analysis in triple NFI KO and control cells shows reduction of the NFI signal on Greek Islands (Supplemental Figure S7F). The lack of consensus NFI motifs from these enhancers, however, and our inability to detect NFI binding on Greek Islands by crosslinked ChIPs (data not shown), suggests that the CUT&RUN signal may reflect indirect recruitment of these factors via protein-protein interactions with Lhx2/Ebf/Ldb1. Regardless of the mechanism of potential NFI binding on OR promoter and enhancers, our data show that NFI proteins establish chromatin modification and nuclear organization landscapes that repress zone 2/3 ORs and enable zone 4/5 ORs in ventral OSNs.

## Discussion

We uncovered the mechanism by which a random process becomes skewed, transforming the relative position of a cell across the dorsoventral axis of the MOE into biased OR gene choices. The solution to the perplexing segmentation of the MOE into overlapping zones is remarkably simple: a gradual increase of heterochromatin deposition and genomic compartmentalization from dorsal to ventral zones, alters, in a sequential fashion, the OR indexes that can be activated in each OSN. ORs that are activatable in one zone, will be silenced in more ventral ones, while ORs that lack heterochromatin but cannot interact with the Greek Island hub in this zone, acquire this privilege in more ventral segments. This serial progression is conceptually reminiscent of the fascinating phenomenon of spatial collinearity in Hox gene regulation(Lewis, 1978; Noordermeer and Duboule, 2013; Noordermeer et al., 2011), albeit with two important distinctions: First, in Hox gene regulation, continuous binding of PcG proteins on Hox clusters is required to maintain long-term gene silencing(Beuchle et al., 2001; Coleman and Struhl, 2017), whereas weeks, or even months, after NFIs are deleted in mOSNs, zonal OR expression patterns are faithfully preserved. Although without a doubt the post-mitotic nature of mOSNs contributes to this stability, it is quite striking that both nuclear architecture and heterochromatinization are preserved in the absence of the sequence-specific factors that initiated their assembly. Second, the order of Hox gene activation is determined by their linear order on the chromosome, while here this order is defined by the sequential OR recruitment in 3-dimensional genomic compartments. Consequently, whereas Hox transcription is highly stereotypic within segments, the activation of an OR allele with zonal-appropriate credentials is not. As our Dip-C analysis revealed, OR compartments have highly variable constitution, with the 3D maps of OR contacts being different even between OSNs from the same zone. This immediately suggests that OR compartments and Greek Island hubs assemble in an “opportunistic” fashion, influenced by both the relative affinities between OR loci and their proximity in the nuclear space. While the former is determined by the position of the OSN, the latter is random and dependent on the relative position of each chromosome during the exit of the last mitosis. With OR gene clusters being located in 18 different chromosomes, it is not surprising that the constitution of OR compartments and Greek Island hubs differs from cell to cell. Thus, although a mechanism evolved to establish some semblance of order in OR gene regulation, the concept of transcriptional activation via interchromosomal associations renders chance a key operating principle.

### A quantitative model for opposing heterochromatin functions

The focus of our experiments was to determine the mechanism that biases OR gene expression to a specific zone. We showed that the concentration of Nfi proteins in OSN progenitors determine the repertoire of ORs that can be expressed in mOSNs via regulation of heterochromatin assembly and nuclear architecture. A possible explanation of the homeotic transformations occurring upon NFI deletion is that different levels of H3K9me3/H3K79me3 enrichment have different consequences in chromatin compaction, genomic compartmentalization, and transcriptional activation. We propose that high levels of H3K9me3/H3K79me3 induce both chromatin compaction and formation of OR compartments; intermediate levels of H3K9me3/H3K79me3 suffice for OR compartmentalization but not for chromatin compaction, while low levels of these modifications do not promote either (Figure 8). Consequently, in each OSN ORs from earlier zonal indexes are not expressed because they are actively silenced by constitutive heterochromatin and ORs from subsequent zones are not expressed because they cannot associate with other ORs and the Greek Island hub (Figure 8). Presumably, the repressive histone modification H3K9me2, which decorates ORs even in other tissues(Magklara et al., 2011), is sufficient to keep the ventral ORs transcriptionally silent in the absence of interactions with the multi-enhancer hub. Thus, only ORs from the correct zonal index combine a partially accessible chromatin state with the ability to associate with the Greek Island hub, restricting OR gene choice to a specific OR repertoire.

**Figure 8.**
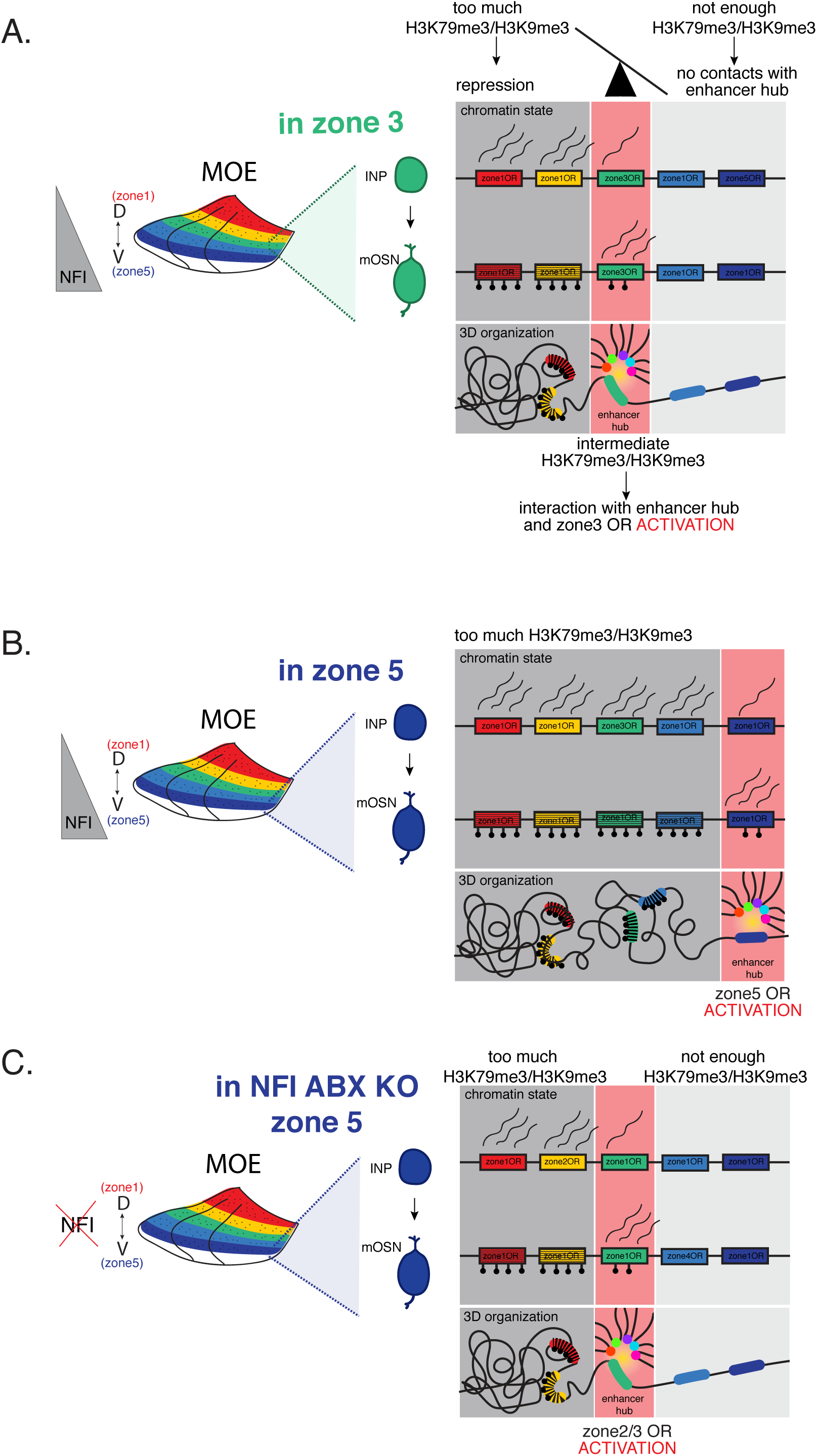
(shall become graphical abstract): NFI-dependent increase of heterochromatin and genomic compartmentalization across the dorsoventral axis determines which OR indexes can be chosen in each zone. (A) In zone 3, during the INP to OSN transition there is higher transcription of zone 1-2 OR indexes than zone 3 OR indexes, resulting in high levels of H3K9me3/H3K79me3 in zone 1-2 ORs, intermediate levels in zone 3 ORs and low levels in zone 4-5 ORs. Consequently zone 1-2 ORs are transcriptionally silenced, while zone 3 ORs have access to the Greek Island hub, and thus can be chosen for singular expression. Zone 4-5 ORs are devoid of both H3K9me3/H3K79me3 and long-range contacts with other ORs and Greek Islands and thus cannot be chosen. Ubiquitous H3K9me2 is likely to keep these ORs completely silent in the absence of activating genomic interactions. (B) Similar with A, but in zone 5, where NFI levels are highest, everything has shifted ventrally. Thus, ORs 1-4 all have high levels of H3K9me3/H3K79me3 and only zone 5 ORs have intermediate levels compatible with interactions with the Greek Island hub and singular activation in mOSNs. (C) Triple deletion of NFI paralogues reverts heterochromatin distribution and genomic compartmentalization to the landscape of zone 3, resulting in ectopic zone 3 OR expression. Similarly, zone 4 NFI KO OSNs may revert to zone 2 landscapes, resulting in ectopic zone 2 OR expression.

The aforementioned model assumes that H3K9me3/H3K79me3 have independent functions in the regulation of chromatin compaction and 3D organization. The former likely depends on high density deposition of tri-methyl marks across numerous successive nucleosomes(Canzio et al., 2011; Canzio et al., 2013). This would allow Hp1 proteins and other components of heterochromatin to polymerize on the chromatin fiber, insulating regulatory sequences from transcriptional activators. The latter, which may be organized by homotypic chromatin contacts(Gibson et al., 2019; Narlikar, 2020) and the phase separation properties of heterochromatin(Mirny et al., 2019), could be accomplished at intermediate levels of enrichment that do not induce chromatin compaction. Consistent with this, recent reports provide functional evidence for a role of H3K79me3/me2 in the establishment of long-range enhancer-promoter interactions(Godfrey et al., 2020) and role of H3K9me3 in establishment of genomic compartments *in vivo* (Falk et al., 2019). Importantly, the differences in H3K9me3/H3K79me3 enrichment and frequency of Hi-C contacts are strongest in iOSNs during the process of OR gene choice. This suggests that there is a transient “bivalent” chromatin modification state during which converging ORs with intermediate levels of heterochromatin can either become fully heterochromatic and transcriptionally silent or associated with Greek Islands and transcriptionally chosen.

In addition to the putative role of histone modifications in the establishment of repressive or activating genomic interactions, the repertoire of transcription factors bound on OR promoters and enhancers could also influence genomic compartmentalization. Increased abundance of NFI motifs on zone 4/5 OR promoters and detection of NFI proteins on Greek Islands by CUTt&RUN are consistent with an NFI-dependent re-direction of the Greek Island hubs towards zone 4/5 ORs though homotypic interactions. Such homotypic interactions by DNA-bound transcription factors were recently shown to promote assembly of specific interchromosomal compartments in yeast(Kim et al., 2019; Kim and Shendure, 2019). That said, we have not demonstrated binding of NFI proteins on zone 4/5 OR promoters and we cannot unequivocally exclude that NFI occupancy of Greek Islands is a reflection of their highly accessible state, since these sequences do not contain *bona fide* NFI motifs. Thus, indirect mechanisms by which NFI expression influences nuclear architecture and OR gene choice cannot be excluded, especially in light of the numerous other zonal transcription factors identified by our experiments. That said, the three NFI paralogues are essential for transforming cellular locations along the dorsoventral MOE axis into zonal OR expression patterns.

Finally, an alternative model for our genomic and genetic observations is that the role of Nfi proteins is to strictly to repress expression of zone 2/3 ORs in zone 4/5 OSNs. In this vein, both marking by H3K9me3/H3K79me3 and recruitment to OR compartments are exclusively repressive, eliminating the competitive advantage that more dorsal OR promoters may have over the promoters of zone 4/5 ORs. Consequently, in a wild type ventral OSN the Greek Island hub is forced to associate with a zone 4/5 OR because the other ORs are inaccessible. In contrast, in the triple NFI KO silencing is alleviated from a fraction of zone 2/3 ORs, which prevail over the zone 4/5 ORs. Thus, sequential silencing of the strongest available OR promoters along the dorsoventral axis is an equally appealing mechanism for MOE segmentation. However, the observation that ectopically expressed zone 3 ORs have higher levels of H3K79me3 in the triple NFI KO than the zone 3 ORs that are not ectopically expressed, supports the notion that an “optimal” level of heterochromatic marks and genomic interactions determines the privileged ORs that can be chosen in each OSN. This would also explain the counterintuitive observation that deletion of H3K9 methyltransferases G9a/Glp abrogates the expression of >99% of OR genes(Lyons et al., 2014).

### Transcription (factor) mediated silencing as a means of establishing zonal restrictions

An intriguing observation from our experiments is that the first detection of H3K9me3/H3K79me3 on OR genes coincides with the onset of high levels of OR transcription at the INP to iOSN transition. Moreover, at the onset of heterochromatin assembly, OR genes have a stronger signal downstream of the OR TSS that declines after the transcription end site (TES), whereas OR enhancers have symmetrical flanking signal, consistent with a bidirectional transcription pattern of distant enhancers (Mikhaylichenko et al., 2018; Wang et al., 2011). These observations are consistent with a mechanism of transcription-mediated heterochromatin assembly on OR genes and their enhancers, with the former being biased towards the gene body and the latter being bi-directional. Biochemical experiments reporting purification of mammalian elongating RNA polymerase II complexes containing Dot1l, the H3K79 lysine methyltransferase (Kim et al., 2012) are also supportive of this hypothesis. Similarly, transcription-dependent deposition of H3K9me2/3 has been observed in *Sacharomyces pompe* and *Arabidopsis thaliana(Allshire and Madhani, 2018)*, providing precedent to the counterintuitive concept of transcription mediated silencing. Alternatively, it is possible that transcription factor binding on OR promoters and enhancers determines directly the levels of H3K9me3/H3K79me3 at the surrounding sequences, explaining why Greek Islands, which are strongly bound by Lhx2, Ebf and likely Nfi, have the highest flanking levels of heterochromatin. In this vein, zone 1 OR promoters, which have the strongest ATAC signal in each zone, also have the strongest levels of H3K9me3/H3K79me3 in each zone.

If relative promoter strength determines directly, or through a transcription-mediated deposition, the levels of heterochromatic marks on each OR, then why do these levels increase for every OR ventrally? Considering that Ebf and Lhx2 expression levels are homogeneous across zones, additional factors have to be deployed to strengthen OR promoter activity in more ventral zones. Thus, we propose that zone 1 promoters are strong enough to induce low transcription and intermediate levels of H3K79me3/H3K9me3 in zone 1 INP3 cells. In subsequent zones, binding of auxiliary factors on zone 1 promoters would further increase their strength, resulting in increased deposition of heterochromatic marks, and resulting in their silencing. Similar principles may apply for all the other OR indexes, with zone 5 promoters being the weakest of all, and thus requiring the highest levels of NFI activity to overcome the lack of strong Ebf motifs. A direct prediction of this hypothesis is that reducing the strength of a zone 1 OR promoter would shift the expression of this OR to more ventral zones. In fact, this is exactly what was reported for the targeted mutation of a pair of Ebf and Lhx2 binding sites in the promoter of Olfr151, which results in shifting the expression of this zone 1 OR to zones 2/3(Rothman et al., 2005).

Importantly, there are additional genetic observations supporting a model of increasing OR promoter strength along the dorsoventral axis. Zone 5 ORs Olfr1507-1509 cannot be expressed as transgenes without the H enhancer(Serizawa et al., 2005), while zone 1 and 2 ORs can be expressed as minigenes with minimal promoter sequences(Vassalli et al., 2002) (and Ben Shykind personal communication). Furthermore, deletion of the H enhancer from the endogenous cluster impairs only the expression of zone 5 and not of zone 1 ORs(Fuss et al., 2007; Nishizumi et al., 2007), supporting the notion that OR promoter strength decreases in ventral OR indexes. Finally, the increased frequency of Greek Island interactions and the somewhat higher complexity of Greek Island hubs in zone 4/5 compared to zone 1, could represent a compensatory mechanism for the weakening of ventral OR promoters. Thus, although OR gene choice may be fully dependent on interactions with the Greek Island hub, ORs from dorsal zones, due to sufficient promoter strength, may be less dependent on *cis* enhancers for the early transcriptional process that facilitates the recruitment to the hub.

### Zonal OR expression preserves a wide receptive field

The final question we wish to address regards the potential benefit of this unusual regulatory mechanism and the merit of generating zonal boundaries in the olfactory epithelium. It is well established that the zonal topography of OR expression in the MOE is reflected in targeting of OSN axons to the olfactory bulb. Because this topography disintegrates in higher olfactory processing centers(Stettler and Axel, 2009), we propose that the major advantages of zonal OR expression relate to the process of OSN axon guidance, the singular nature of OR expression in mature OSNs, and the distributed representation of every OR identity. Containment of OSNs expressing the same OR at defined anatomical segments of the MOE, facilitates the fasciculation of their axons and reduces the number of different extracellular identities required for the formation of OR-specific glomeruli. In other words, there is no need for 1400 extracellular barcodes for the generation of unique glomeruli, but only 40-200 would be sufficient for the segregation of the axons emerging from a specific zone. Furthermore, limiting the process of OR gene choice to 40-200 ORs instead of 1400 different ORs, reduces the probability of choosing two different ORs at the same time. Finally, this process provides a safeguard against the evolution of transcriptionally dominant OR alleles. If an OR promoter was to become too strong, it would be chosen first, suppressing the choice of other ORs. However, with this elegant regulatory mechanism in place, stronger promoters induce early transcription in INPs and accumulation of high levels of heterochromatin outside of zone 1, leading to their “ostracization” in zones 2-5. Of course, this system does not have an unlimited capacity to block transcriptional domination, since artificially powerful promoters can express OR transgenes in high OSN frequencies(D’Hulst et al., 2016; Fleischmann et al., 2008; Nguyen et al., 2007). On the other hand, weaker promoters can still be represented at considerable frequencies, since there are zones dedicated to promoters of various strengths, with zone 5 likely preserving frequent expression of the 40 weakest OR promoters. Importantly, as eluted by lack of consequences in zonal OR transcription by the “late” NFI deletion, differences in promoter strength may become irrelevant in mOSNs, where accumulation of numerous Ebf/Lhx2/Ldb1 bound enhancers assembles a transcriptional complex that supports high transcriptional levels for every chosen OR. Thus, by combining developmental restrictions to the OR identities that can be chosen in each OSN, with a random selection from this spatially determined OR repertoire, the olfactory organ expresses thousands of OR alleles at meaningful frequencies, assuring sensitive detection of the highest number of volatile chemicals.

## Acknowledgments

The authors have no competing interests for this work. Mice were treated in compliance with the rules and regulations of IACUC under protocol number AC-AAAT2450 and AC-AABG6553. We thank members of the Lomvardas lab for critical discussions and input on the manuscript. We thank Drs. Tom Maniatis, Richard Axel, Gary Struhl, and Abbas Rizvi for critical comments and discussions. Stavros Lomvardas acknowledges support from the National Institutes of Health Common Fund 4D Nucleome Program (Grant 5U01DA040582), and the National Institute of Deafness and Communications Disorders (Grant 5R01DC018745). SL was also supported by the HHMI Faculty Scholar Award (#55108540). RMG was supported by NYSTEM contracts C030133 and C30290GG. GB was supported by 5R01DC013561 (NIH).

## Data Availability

All of the sequencing data (RNA-seq, scRNA-seq, ChIP-seq, Omni-ATAC, CUT&RUN, HiC and Dip-C) reported in this paper are publicly available at GEO under accession number_______. HiC and Dip-C data can also be accessed at the 4DN Data Portal (https://data.4dnucleome.org/). This paper also makes use of published Olfr17+ mOSN Hi-C data available at the 4DN Data Portal under the accession number 4DNESNYBDSLY.

## STAR Methods

### EXPERIMENTAL MODEL AND SUBJECT DETAILS

Mice were treated in compliance with the rules and regulations of IACUC under protocol number AC-AAAT2450 and AC-AABG6553. Mice were sacrificed using CO2 following cervical dislocation. A complete list of mouse genotypes used for every experiment is in the Table2. Mash1-CreER (also known as Ascl1^CreERT2^)(Kim et al., 2011); Ngn1-GFP(Magklara et al., 2011) and Cre inducible tdTomato reporter (also known as *B6N.129S6-Gt(ROSA)26Sor^tm1(CAG-tdTomato*,-EGFP*)Ees/J^*)(Madisen et al., 2010) mice were used to isolate four cell types in the olfactory lineage (GBC: tdTomato+ GFP-, INP: tdTomato+ GFP+, iOSN: tdTomato-GFP+ (bright), and mOSN: tdTomato+ GFP dim) by sorting cells 48 hours after tamoxifen injection. GFP bright and dim cells from Ngn1-GFP pups (P6) were also used to isolate a mix of INP/iOSN cells and mOSN cells respectively. Omp-ires-GFP(Shykind et al., 2004) mice were used to isolate mature OSNs from adult (> 8-week-old) mice. In order to obtain zonal iOSNs and mOSNs, Olfr1507-ires-Cre(Shykind et al., 2004) and tdTomato alleles were crossed in with either Ngn1-GFP or Omp-ires-GFP alleles to aid in zonal dissection (by labeling Ollfr1507+ expressing cells in zone5).

Early knockout of Nfi A, B, and X (NfiABX) in horizontal basal cells (HBSs: the stem cell of the olfactory epithelium) was achieved by crossing NfiA fl/fl NfiB fl/fl and NfiX fl/fl triple conditional alleles, described in (Clark et al., 2019), with KRT5-CreER(Rock et al., 2009) and tdTomato. Adult mice (> 8-week-old) had deletion of NfiABX in horizontal basal cells induced with 3 intraperitoneal injections with tamoxifen (24 hours apart). Ten days after the first injection, the olfactory epithelium was ablated with one intraperitoneal injection of methimazole, inducing proliferation of the HBCs and regeneration of a Nfi ABX knockout olfactory epithelium. The olfactory epithelium was allowed to regenerate for 40 days before collecting the MOE and FAC-sorting the tdTomato+(dim) cell population, which contains a mixture of mostly mOSNs and some INP and iOSN cells, as described in detail in (Monahan et al., 2019). For some experiments Omp-ires-GFP was crossed in to ensure all cells collected were mOSNs. Late knockout of Nfi ABX in mOSNs was achieved by crossing NfiA, B, and X triple conditional alleles with tdTomato and Omp-ires-Cre, and FAC-sorting tdTomato+ cells from adult mice. Complete list of all the mouse genotypes can be found in Table 2.

### METHOD DETAILS

#### Zonal Annotation

OR genes were assigned a zonal annotation (referring to their native zone of expression) based on (Tan and Xie, 2018). We generated bins from the continuous data by rounding to the nearest integer. There are a total of 1011 ORs with known zonal annotation. Of these, 115 are ClassI ORs, of which nearly all are expressed in zone1, and 896 are ClassII ORs, of which 261 are expressed in zone1, 283 in zone2, 164 in zone3, 144 in zone4 and 44 in zone5.

Zonal dissection of the olfactory epithelium:

We used the fluorescent signal in Olfr545-delete-YFP(Bozza et al., 2009) (zone 1 OR), Olfr17-ires-GFP (Shykind et al., 2004)(zone 2 OR), and Olfr1507-ires-GFP(Shykind et al., 2004) (zone 5 OR) mice to practice dissections of zones 1, 2/3, and 4/5, respectively. Upon obtaining an accurate understanding of the zonal boundaries in the MOE we performed zonal dissections without the use of these fiduciary markers. Accuracy of dissections was confirmed by RNA-seq. For some experiments Olfr1507-ires-Cre and tdTomato reporter was crossed in to assist with accurate zone5 dissection (see Table 2.)

#### Fluorescence-activated cell sorting

Cells were prepared for FAC-sorting as previously described in(Monahan et al., 2019) by dissociating olfactory epithelium tissue with papain for 40 minutes at 37°C according to the Worthington Papain Dissociation System. Cells were washed 2x with cold PBS before passing through a 40-um strainer. Live (DAPI-negative) fluorescent cells were collected for RNA-seq, native ChIP-seq, omni-ATAC-seq and CUT&RUN. Alternatively, for Hi-C cells were fixed for 10 minutes in 1% formaldehyde in PBS at room temperature, quenched with glycine, and washed with cold PBS before sorting fluorescent cells. Alternatively, for Dip-C, cells were fixed in 2% formaldehyde in PBS at room temperature for 10 minutes, inactivated with 1% BSA, and washed with cold 1% BSA in PBS before sorting fluorescent cells. All cells were sorted on a BD Aria II.

#### Single cell RNA-seq in olfactory lineage cell types

Mash1-CreER; tdTomato; Ngn1-GFP pups (ages P2-P4) were injected with tamoxifen and olfactory epithelium was collected after 48 hours. The tissue was dissected into ventral (zone 3-5) and dorsal OE (zone1-2) sections, from which GBC (tdTomato+, GFP-), INP (tdTomato+, GFP+), iOSN (tdTomato-, GFP+ bright) and mOSN (tdTomato-, GFP dim) cells were sorted into 384 well plates (split between the cell types). Each well of the 384 well plate had unique cell and molecular barcodes. Library preparation and sequencing was performed in collaboration with the New York Genome Center (NYGC) using a TSO approach for library preparation and sequenced on HiSeq2500. Reads were aligned to the mm10 genome according to the Drop-seq(Macosko et al., 2015) pipeline (http://mccarrolllab.org/dropseq/), which uses STAR for alignment, and discarding multi mapped reads with Samtools -q 255. Aligned single cells had a median of 133,686 unique transcripts (UMIs) and 2,331 genes per cell (detected with a threshold of at least 3UMI). Experiment was performed in biological replicate, resulting in 764 cells, from which we discarded cells with less than 1000 genes and 20,000 UMIs, resulting in 669 cells. We further filtered for cells that contained less than 5% mitochondrial reads, resulting in 591 cells used for analysis. We used Seurat v3 to normalize counts and cluster single cells, resulting in 5 populations. Clusters were assigned a cell-type based on expression of known olfactory lineage markers. We used the default setting of genes expressed in at least 3 cells for clustering but changed it to 1 when looking at OR expression (since expression of any OR out of > 1000 genes is a rare event). For all OR expression analysis we used a threshold of 3UMI for an OR to be considered expressed.

#### Bulk RNAseq in olfactory lineage cell types

GBC, INP, iOSN and mOSN were isolated from Mash1-CreER; tdTomato; Ngn1-GFP pups as described above with the tissue being dissected into a ventral (mostly zone 4-5), dorsal OE (mostly zone1) and a central section (that is enriched for zone2-3). The experiment was performed in biological replicate. RNA was extracted from FAC-sorted cells using Trizol and libraries were prepared with Nugen NuQuant RNA-seq library system and sequenced 50PE on HiSeq2500 or 75PE NextSeq (and trimmed to 50bp before aligning). Cutadapt was used to remove adapter sequences and reads were aligned to the mm10 genome with STAR. Samtools was used to select high mapping quality reads (-q 30). Normalization, calculation of FPKM (which we converted to TMP), and differential expression analysis was performed in R with DEseq2. For all RNA-seq data p-values refer to adjusted p-value (padj) calculated in DEseq2.

To find zone5 enriched transcription factors at each developmental stage we determined significantly differentially expressed transcription factors (from the Gene Ontology database annotation “DNA binding transcription factor activity”) between zone4/5 and zone1 cells with a padj less than 0.05 and at least a twofold change in expression (see Table 1.) To get the most likely candidates driving zonal identity we further filtered the list for TFs with at least a 3-fold difference between zone 1 and zone 4/5 cells, and an expression level of at least 15 TPM).

#### Zonal vs non-zonal mOSN markers from olfactory lineage RNA-seq data

To find zone4/5 enriched mOSN markers, we looked at non-OR genes differentially expressed between zone4/5 and zone1 or zone2-3 mOSNs (tomato-, GFP dim cells) with padj less than 0.05, and at least a twofold change in expression, of which there were 208; and performed the inverse analysis to generate a list of z1 or z2 enriched mOSN markers, of which there were 141 genes. To find non-zonal mOSN markers, we made a list of significantly upregulated genes (with a padj less than 0.05, and a fold change greater two) in mOSNs (tomato-, GFP dim cells) across all zones compared to iOSNs (tomato-, GFP+ bright cells) across all zones. We further filtered out genes that were significantly differentially expressed between zone5 and z1 or 2 mOSNs and took the top 200 most significant genes.

#### RNA-seq in Nfi knockout zone5 mOSNs

To look at gene expression changes resulting from Nfi deletion in olfactory progenitors we used Nfi triple knockout (NFI A,B,X fl/fl, tdTomato, OMP-gfp, Krt5-CreER), AB only double knockout (NFI A,B fl/fl, tdTomato, OMP-gfp, Krt5-CreER), X only knockout (NFI X fl/fl, tdTomato, OMP-gfp, Krt5-CreER) or wt (tdTomato, OMP-gfp, Krt5-CreER) mice and followed the induction protocol for early knockout, described above. After rebuilding the MOE from knockout progenitors, we dissected zone4/5 and FAC-sorted GFP+ mOSNs. RNA was extracted from sorted cells using Trizol and RNA-seq libraries were prepared with Nugen Nuquant RNA-seq library prep kit and sequenced 75PE on Nextseq 550. Reads were aligned exactly as described for zonal olfactory lineage data and similarly DEseq2 was used to determine differentially expressed genes between the different knockout and wt cells. To determine if zone4/5 mOSN, zone1/2 mOSN and non-zonal mOSN markers change in Nfi knockout zone5 we looked at expression differences we analyzed the expression differences of the genes our markers lists.

#### Native chromatin immunoprecipitation from FAC-sorted cells

Native chip was performed as described in detail (Monahan et al., 2017). Unless otherwise indicated all steps were carried out at 4°C. Briefly, FAC-sorted cells were pelleted at 600 rcf for 10 minutes in a swinging bucket centrifuge at 4°C and resuspended in cold Buffer I (0.3M Sucrose, 60 mM KCl, 15 mM NaCl, 5 mM MgCl2, 0.1 mM EGTA, 15 mM Tris-HCl pH 7.5, 0.1 mM PMSF, 0.5 mM DTT, 1x protease inhibitors). Cells were lysed by adding equal volume cold BufferII (Buffer I with 0.4% NP40) and incubating for 10 minutes on ice. Nuclei were pelleted 10 min at 1000 rcf and resuspended in 250ul cold MNase buffer (0.32M Sucrose, 4 mM MgCl2, 1 mM CaCl2, 50 mM Tris-HCl pH 7.5, 0.1 mM PMSF, 1x protease inhibitors). Micrococcal Nuclease digestion was carried out by adding 0.1U Micrococcal Nuclease (Sigma) per 100ul buffer and incubating for 1 min 40 sec in a 37°C water bath, then stopping the digestion by adding EDTA to a final concentration of 20mM. The first soluble chromatin fraction (S1) was collected by pelleting nuclei 10 min at 10,000 rcf at 4°C and taking the supernatant to store at 4°C overnight. Undigested, pelleted material was resuspended in 250ul cold Dialysis Buffer (1 mM Tris-HCl pH 7.5, 0.2 mM EDTA, 0.1 mM PMSF, 1x protease inhibitors) and rotating overnight at 4°C. The second soluble chromatin fraction (S2) was collected by pelleting insoluble material 10 min at 10,000 rcf at 4°C and taking the supernatant. S1 and S2 chromatin fractions were combined and used for immunoprecipitation with 5% material being retained for input. Equal cell numbers were used for control and knockout IPs, or between different cell types or zones. To perform immunoprecipitation (IP), chromatin was diluted to 1ml in Wash Buffer1 (50mM Tris-HCl pH 7.5, 10 mM EDTA, 125 mM NaCl, 0.1% Tween-20, 5 mM 1x protease inhibitors) and rotated overnight at 4°C with 1ug antibody. Dynabeads (10ul Protein A and 10ul Protein G per IP) were blocked overnight with 2 mg/ml yeast tRNA and 2 mg/mL BSA in Wash Buffer 1. Blocked beads were washed once with Wash Buffer 1, then added to antibody bound chromatin and rotated 2-3 hours at 4°C. Chromatin bound beads were washed 4x with Wash Buffer1, 3x with Wash Buffer 2 (50mM Tris-HCl pH 7.5, 10 mM EDTA, 175 mM NaCl, 0.1% NP40, 1x protease inhibitors), and 1x in TE pH 7.5. IP’d DNA was eluted by resuspending beads in 100 uL Native ChIP Elution Buffer (10 mM Tris-HCl pH7.5, 1 mM EDTA, 1% SDS, 0.1 M NaHCO3) in a thermomixer set to 37°C and 900 rpm for 15 minutes, repeating the elution 2x and combining the eluates. IPs and inputs (diluted to 200ul in elution buffer) were cleaned up with Zymo ChIP DNA columns (Zymo Research, D5205). Libraries were prepared with NuGEN Ovation V2 DNA-Seq Library Preparation Kit, and sequenced 50PE on HiSeq2500 and or 75PE on NextSeq 550.

#### Native ChIP-seq analysis

Sequenced reads were pre-processed by trimming adapters with Cutadapt, then aligned to the mm10 genome using Bowtie2, with default setting except for maximum insert size set to 1000 (-X 1000), allowing larger fragments to be mapped. Duplicate reads were removed with Picard, and high mapping quality reads were selected with Samtools (-q 30). After confirming replicates looked similar, they were merged with HOMER and used to generate signal tracks at 1bp resolution normalized to a library size of 10,000,000 reads. Signal density over OR genes was calculated with HOMER annotatePeaks.pl then normalized to the length of each OR gene. Native ChIP heatmaps were generated with deeptools with OR gene bodies re-scaled to 6kb and showing 2kb flanking on each side.

#### Omni ATAC-seq

Omni ATAC was performed on 50,000 iOSNs from zone1 or zone4/5 (isolated from Mash1-CreER, tdTomato, Ngn1-GFP mice as described in detail above). The protocol was carried out as described in(Corces et al., 2017) Briefly, FAC-sorted cells were pelleted at 500 rcf at 4°C for 5 min and lysed by resuspended in 50ul cold ATAC-Resuspension Buffer (RSB) (20 mM Tris-HCl pH 7.6, 10 mM MgCl2, 20% Dimethyl Formamide) containing 0.1% NP40, 0.1% Tween-20, and 0.01% Digitonin, pipetting gently 3x, and incubating 3 min on ice, then washing out with 1ml ATAC-RSB containing 0.1% Tween-20 but with no NP40. Nuclei were pelleted at 500 rcf for 10 min at 4°C, and the supernatant was aspirated, making sure to remove every trace from the nuclei pellet. Transposition of chromatin was carried out by resuspending nuclei in 50ul cold transposition mixture (25 ul 2x TD buffer, 2.5 ul Tn5 transposase (100nM final), 16.5 ul PBS, 0.5 ul 1% digitonin, 0.5 ul 10% Tween-20, 5ul water) and incubating in a thermomixer set to at 37°C and 1000 rpm mixing for 30 minutes. Transposed fragments were cleaned up with Zymo DNA Clean and Concentrator-5 Kit and eluted in 21 ul elution buffer. Library was prepared by amplifying DNA initially for 5 cycles using 2x NEBNext MasterMix (and adding adaptors with a unique barcode to each sample), then running a qPCR to determine additional cycles. After final amplification libraries are cleaned up with Zymo DNA Clean and Concentrator-5 Kit. Experiment was performed in biological replicate. Omni-ATAC-seq libraries were sequenced 75PE on Nextseq 550 and reads were pre-processed and aligned to the mm10 genome as described for native ChIP-seq.

#### Omni ATAC-seq analysis

Omni ATAC-seq peaks were called with HOMER using -factor -size 150 -fragLength 150 parameters. Differential peaks were determined with Diffbind after recentering peaks and resizing to to 300bp (summits = 150). There were 14,169 peaks differentially expressed between zone1 and zone 4/5 iOSNs (with an FDR less than 0.05). We used HOMER to identify ne-novo motifs in these differentially accessible peaks.

#### CUT&RUN

CUT&RUN was performed as described in(Skene et al., 2018). After FAC-sorting, cells were pelleted and washed 2x with Wash Buffer, before binding to activated Concanavalin A beads by incubating 10 minutes at room temperature with rotation. 100,000-250,000 cells were used per antibody. Supernatant was removed and beads were resuspended in 200ul Antibody buffer with NIFA or control (rabbit-anti-rat) antibodies at 1:200 dilution and incubated overnight at 4°C with gentle rotation. Beads were washed once with Digitonin Buffer, and Protein A-MNase fusion protein was added at 700ng/ml in digitonin buffer and incubated for 1 hour at 4°C with gentle rotation. Beads were washed 3x in Digitonin buffer and then resuspended in 150ul digitonin buffer and set to cool in a heat block set in an ice bath for ∼ 5 minutes. MNase cleavage was activated with the addition of CaCl2 (2mM final concentration) and placing the tube back in the ice bath to incubate for 5 minutes. Digestion was stopped with the addition of STOP buffer and incubated for 30 minutes at 37°C to release the fragments. Libraries were prepared with NuGEN Ovation V2 DNA-Seq Library Preparation Kit, and sequenced 75PE on NextSeq 550. Reads were pre-processed and aligned to the mm10 genome as described for native ChIP-seq

#### In situ Hi-C

In situ Hi-C and library preparation was performed as exactly as described(Monahan et al., 2019). Briefly, FAC-sorted cells (inputs ranged from 150,000 to 500,000 cells) were pelleted at 500 rcf for 10 minutes and lysed in Lysis buffer (50 mM Tris pH 7.5 0.5% NP40, 0.25% sodium deoxychloate 0.1% SDS, 150 mM NaCl and 1x protease inhibitors) by rotating for 20 min at 4°C. Nuclei were pelleted at 2500 rcf, permabilized in 0.05% SDS for 20 min at 62 °C, then quenched in 1.1% Triton-X100 for 10 min at 37 °C. Nuclei were then digested with DpnII (6U/ul) in 1× DpnII buffer overnight at 37 °C. In the morning, nuclei were pelleted at 2,500*g* for 5 min and buffers and fresh DpnII enzyme were replenished to their original concentration and nuclei were digested for 2 additional hours. Restriction enzyme was inactivated by incubating 20 minutes at 62 °C. Digested ends were filled in for 1.5 hours at 37 °C using biotinylated dGTP. Ligation was performed for 4h at room temperature with rotation. Nuclei were pelleted and sonicated in 10 mM Tris pH 7.5, 1 mM EDTA, 0.25% SDS on a Covaris S220 (16 minutes, 2% duty cycle, 105 intensity, Power 1.8-1.85 W, 200 cycles per burst, max temperature 6°C). DNA was reverse crosslinked with RNAseA and Proteinase K overnight at 65 °C then purified with 2× Ampure beads following the standard protocol and eluted in water. Biotinylated fragments were enriched with Dynabeads MyOne Strepavidin T1 beads and on bead library preparation was carried out with NuGEN Ovation V2 DNA-Seq Library Preparation Kit, with some modifications: instead of heat inactivation following end repair beads were washed 2x for 2 min at 55 °C with Tween Washing Buffer (TWB)(0.05% Tween, 1 M NaCl in TE pH 7.5) and 2x with 10 mM Tris pH 7.5 to remove excess detergent. After ligation of adapters beads were washed 5x with TWB and 2x with 10 mM Tris pH 7.5. Libraries were amplified for 10 cycles and cleaned up with 0.8V Ampure beads. Each experiment was performed with two biological replicates and prepared Hi-C libraries were sequenced 75PE on NextSeq 500.

#### In situ Hi-C analysis

Reads were aligned to the mm10 genome using the distiller pipeline (https://github.com/mirnylab/distiller-nf), uniquely mapped reads (mapq > 30) were retained and duplicate reads discarded. Contacts were then binned into matrices using cooler. (Abdennur and Mirny, 2020). Analysis was performed on data pooled from two biological replicates, after confirming that the results of analysis of individual replicates were similar. Hi-C contact maps of OR clusters on chromosome 2 were generated with raw counts of Hi-C contacts normalized to counts/billion at 100kb resolution. The maximum value on the color scale was set to 150 contacts per 100kb bin. Analysis of zonal OR gene cluster contacts was performed normalized counts binned at 50kb resolution. All analyses were repeated using balanced counts generated by cooler (-mad-max 7), with similar results except balanced matrices discarded almost 10% of OR cluster bins due to relatively poor sequencing coverage.

#### Dip-C

To isolate mature olfactory sensory neurons, Castaneous (Cas) mice were crossed to OMP-ires-GPF mice. MOE was collected from adult heterozygous mice resulting from this cross. The tissue was dissected into zone 1 and zone4/5, fixed for 10 minutes in 2% formaldehyde and FAC-sorted to isolate GFP+ mOSNs. Dip-C was performed as described(Tan et al., 2019) on 96 mature OSNs: 48 each from zones 1 and zones 4/5. Briefly, cells were lysed in Hi-C Lysis Buffer (10mM Tris pH8, 10mM NaCl, 0.2% NP40, 1x protease inhibitors) on ice for 15 minutes, nuclei were pelleted at 2500 rcf for 5 min at 4°C, then resuspended in 0.5% SDS and permeabilized 10 minutes at 62 °C then quenched in 1.1% Triton X-100 15min at 37 °C. Nuclei were digested in 1x DpnII buffer and 6U/ul DpnII enzyme and digested overnight at 37°C. Nuclei were then washed once in Ligation Buffer, and resuspended in Ligation buffer with 10U T4 DNA Ligase (Life Tech), and incubated for 4 hours at 16oC shaking at 600rpm. After ligation nuclei were pelleted and resuspended in cold PBS with DAPI to a final concentration of 300nM and GFP+ cells were FAC-sorted into a 96 well plate with 2ul lysis buffer (20mM Tris pH 8, 20mM NaCl, 0.15% Triton X-100, 25mM DTT, 1mM EDTA, 500nM Carrier ssDNA, and 15ug/mL Qiagen Protease) and lysed for 1 hour at 50°C and inactivated 15 minutes at 70°C. DNA was transposed by adding 8ul transposition buffer (12.5 mM TAPS pH 8.5, 6.25mM MgCl2, 10% PEG 8000) with ∼0.0125 uLTn5 (Vanzyme) and incubated at 55°C for 10 min, then stopped with transposome removal buffer (300nM NaCl, 45 mM EDTA, 0.01% Triton X-100 with 100ug/mL Qiagen Protease) and incubated at 50°C for 40 minutes and 70°C for 20 minutes. Libraries were amplified 14 cycles with i5 and i7 Nextera primers, with unique barcodes for each cell. Libraries from all cells were pooled and cleaned up with Zymo DNA Clean and Concentrate Kit. Libraries were sequenced 150PE on NextSeq 550.

#### Dip-C analysis

Sequenced Dip-C reads were processed according to the Dip-C pipeline (https://github.com/tanlongzhi/dip-c). Reads were aligned to mm10 with BWA mem, and hickit was used to determine the haplotype of each contact based on SNPs between Cas and OMP-ires-GFP mice and make a model of the 3D genome. Since OMP-ires-GFP mice were a mixture of Bl6/129 strains, we only included SNPs that were unique to Cas mice to distinguish homologs. After alignment, cells were filtered using several quality control metrics described in Tan et al. 2019: We excluded cells that had less than 20,000 reads, cells that had a low contact-to-read ratio, and cells that had a high variability in 3D structure across computational replicates. Only 4 of 96 cells failed these metrics. Overall, the median number of contacts across cells was over 400,000. Computational analysis OR genes and Greek Island enhancers, including computing average contact densities and analysis of the 3D models, was performed using the Dip-C pipeline. Average contact densities between OR genes and/or Greek Islands were calculated with ‘dip-c ard’. Pairwise distances between OR genes and/or Greek Islands from the 3D models were extracted with ‘dip-c pd’. Heatmaps of pairwise distance were either ordered by genomic position or reordered using hierarchical clustering. To determine the size of OR gene aggregates and Greek Island hubs, the number of OR genes or Greek Islands within a specified radius of was calculated with ‘network_around.py’. 3D models were visualized with PyMol and used to generate sequential slices of the nucleus.

#### Antibodies

Olfr17 antibody were raised in rabbits against epitope RRIIHRTLGPQKL located at the C-terminus of the OR protein. Olfr1507 antibody was described in (Barnea et al., 2004). NFIA antibody from Active Motif (39397) was used for IF and CUT&RUN. Rabbit-anti-rat antibody from abcam (ab97096brewerb) was used as a control in CUT&RUN. The following antibodies were used for native ChIP: H3K79me3 (abcam ab2621) and H3K9me3 (ab8898).

## Supplemental Figure and Table Legends

**Supplementary Figure S1 (related to Figure 1):**
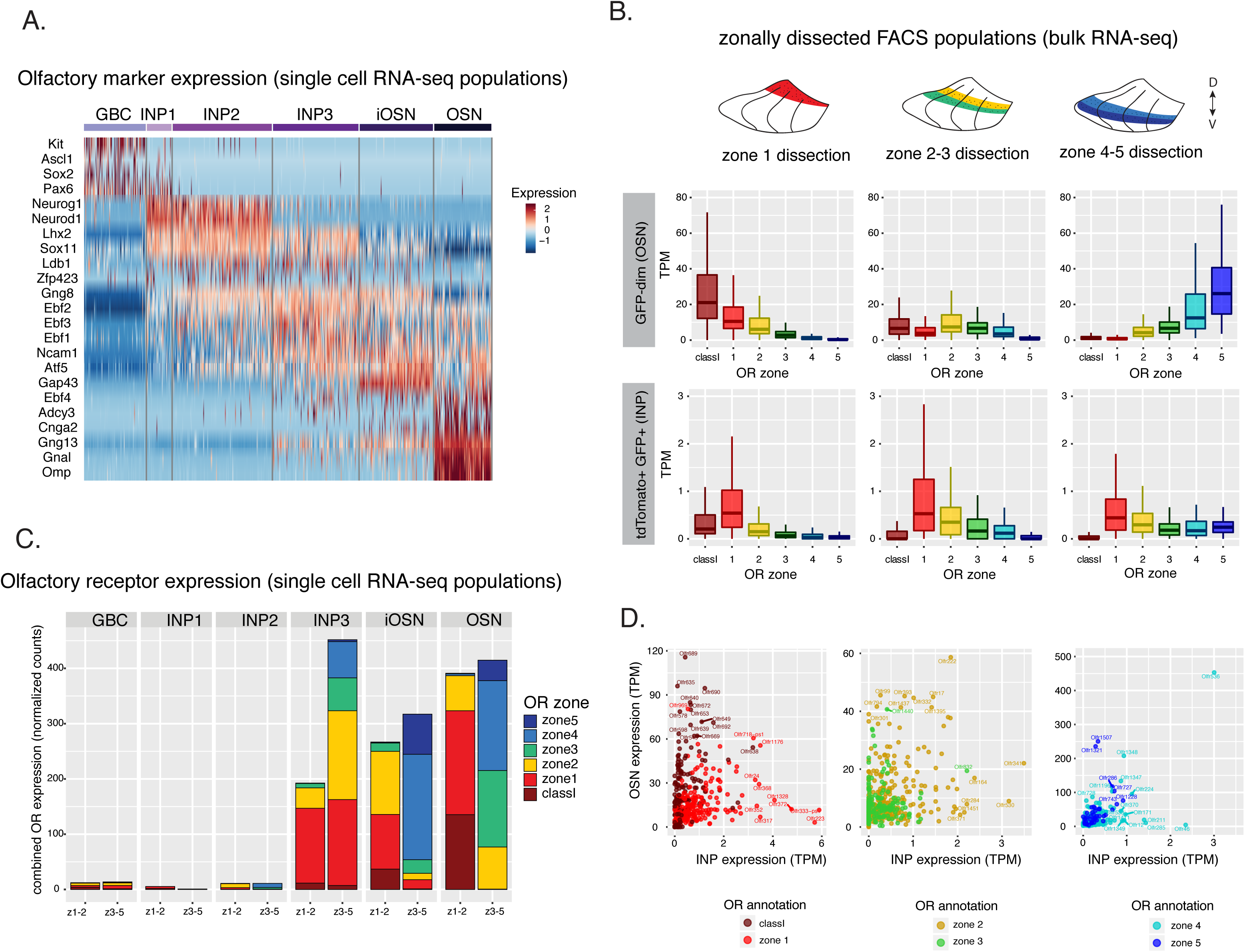
OR genes highly expressed in progenitors are less frequently chosen in mature OSNs. (A) Heatmap shows the expression of known olfactory lineage markers in single cells ordered into columns by their t-SNE cluster identity. Expression is represented in terms of log2 fold change relative to their average expression. (B) Bulk RNA-seq of cell populations isolated using the same experimental strategy outlined in Figure 1A dissected into zone 1 (left) zone 2/3 (middle), and zone 4/5 (right). (C) Aggregate OR expression per zonal cell population from single cell RNA-seq (normalized for the number of cells per zonal population) shows onset of OR expression in INP3 cells and zonal OR preference in different populations. (D) OR expression in INP and mOSN cells shows an anticorrelation between ORs preferentially expressed at the in INP vs mOSN stage.

**Supplementary Figure S2 (related to Figure 2):**
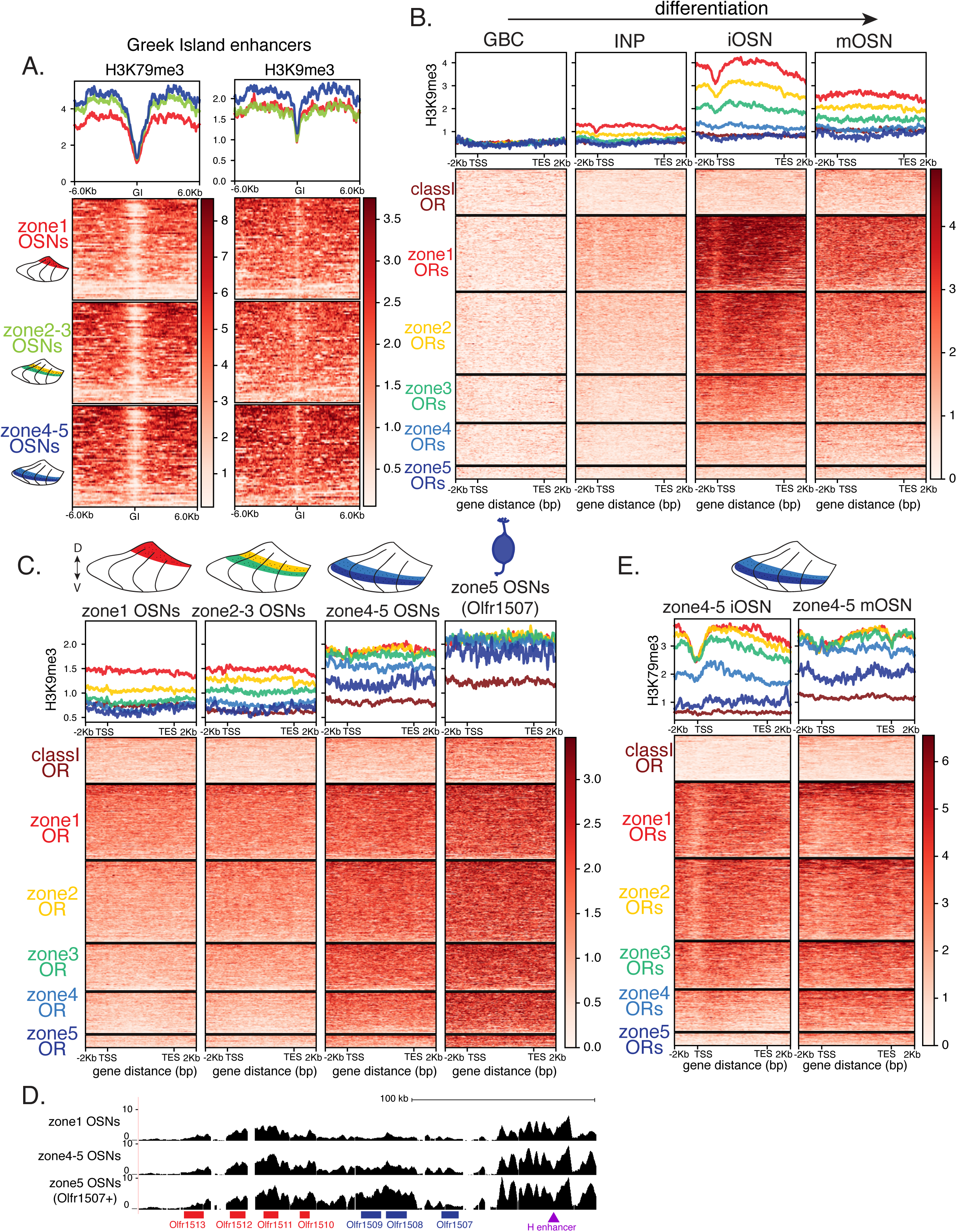
Progressive accumulation of heterochromatin on OR genes occurs in developing OSNs. (A) H3K9me3 and H3K79me3 native ChIP-seq signal over 63 OR enhancers (Greek Islands) in mOSNs from three zonal dissections: zone1 top, zone2/3 middle, and zone 4/5 bottom. Heatmaps centered on Greek Islands show high levels of H3K79me3 and H3K9me3 in mOSNs from all zones. (B) Heatmaps of H3K9me3 native ChIP-seq signal over OR genes in GBC, INP, iOSN and mOSN populations shows a similar onset of deposition to that of H3K79me3. (C) Heatmaps of H3K9me3 native ChIP-seq in mOSNs from dissected zones and a pure population of Olfr1507 (zone 5 OR) expressing mOSNs. (D) Comparison of H3K79me3 native ChIP-seq over OR genes n zone 4/5 iOSNs and mOSNs. H3K79me3 signal is depositred progressively on zone 4/5 ORs as the cells mature. (E) H3K79me3 native ChIP signal track over the OR gene cluster containing Olfr1507 in zone1 mOSNs (top row), zone 4/5 mOSNs (middle row), and Olfr1507-expressing zone5 mOSNs (bottom row). Results show absence of H3K79me3 over the active gene in Olfr1507 expressing mOSNs and near equal signal over all other OR genes regardless of their zone. Purple triangle marks the H Greek Island enhancer.

**Supplementary Figure S3 (related to Figure 3):**
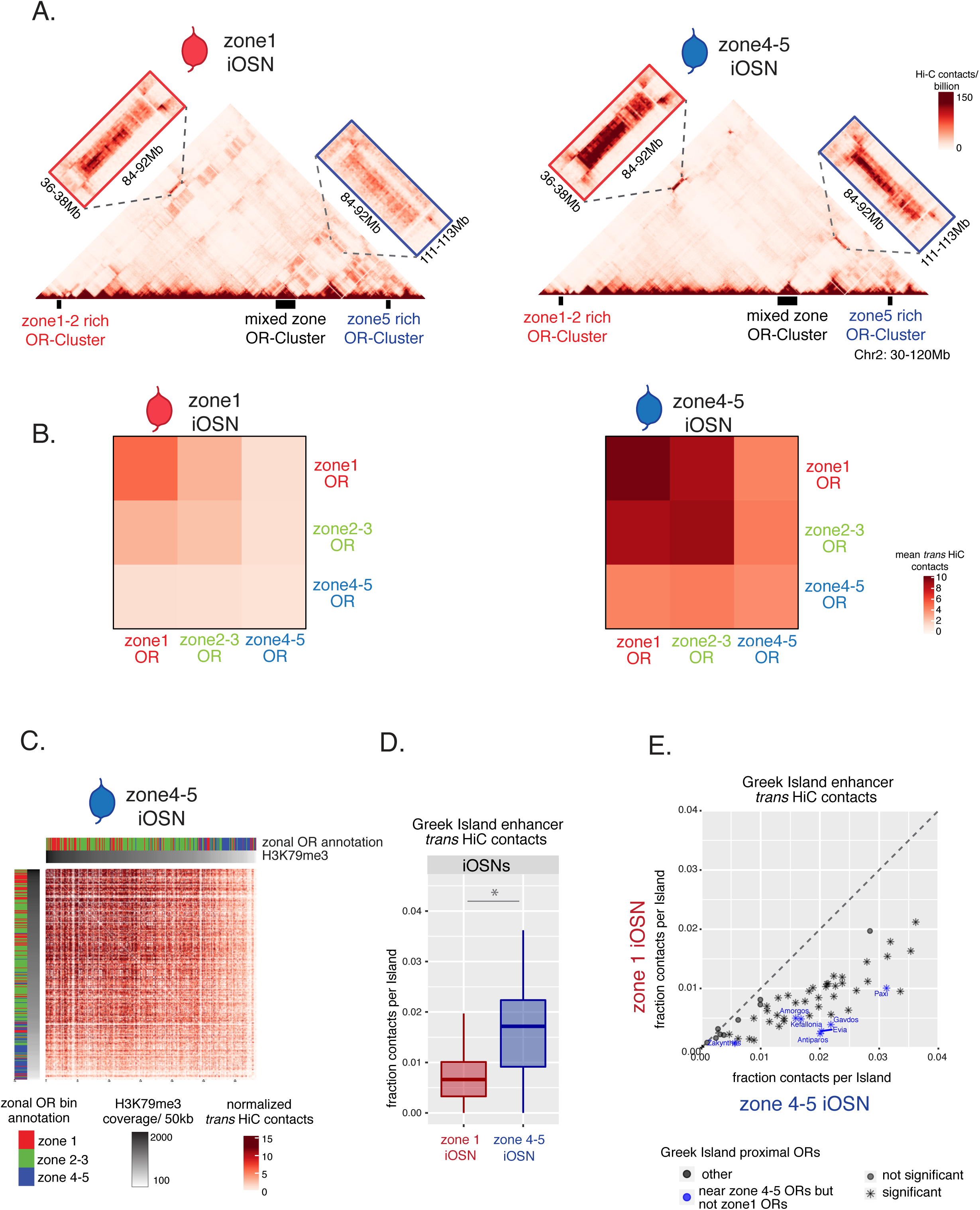
Hi-C in iOSNs shows nuclear structure during OR choice: isolation of ORs from lower zones and increased enhancer interaction. (A) In situ Hi-C contact matrices of a 90Mb region of chromosome 2 in zone1 and zone4/5 iOSNs show long range interactions between three large OR gene clusters. The OR cluster on the right (blue) enriched for zone 4/5 ORs, does not contain any zone1 ORs, and only makes strong interactions with other clusters in zone4 /5 iOSNs. (B) Heatmaps of average interchromosomal HiC contacts between OR genes annotated by their zonal index show increased contacts in zone 4/5 iOSNs compared to zone1 iOSNs. (C) Interchromosomal interactions between OR gene loci are correlated with the amount of H3K79me3 deposition. Heatmap of normalized interchromosomal Hi-C contacts between OR cluster bins. Each bin is annotated according to the zonal index of their resident OR genes: zone1 ORs, red; zone2-3 ORs, green; zone4-5 ORs, blue (shown in the zonal color bar) and ordered by H3K79me3 ChIP signal (gray color bar on top). OR cluster regions that have higher levels of H3K79me3 and are enriched for zone1 ORs have increased *trans* Hi-C contacts. (D) Quantification of interchromosomal interactions between Greek Island enhancers shows they form more frequent interactions in zone4/5 compared to zone1 iOSNs (Wilcoxon rank sum test p-value = 8.12×10-8). (E) Comparison of interchromosomal interactions made by individual Greek Islands shows 47 out of 59 form significantly more contacts in zone 4-5 (Wilcoxon rank sum test: p-value < 0.5). Greek Islands residing in OR clusters with zone 4-5 ORs and no zone1 ORs (blue) are 5 of the top 6 most differentially interacting Greek Islands. All analysis was performed at 50kb resolution and shows merged data from two biological replicates that yielded similar results when analyzed separately.

**Supplementary Figure S4 (related to Figure 4):**
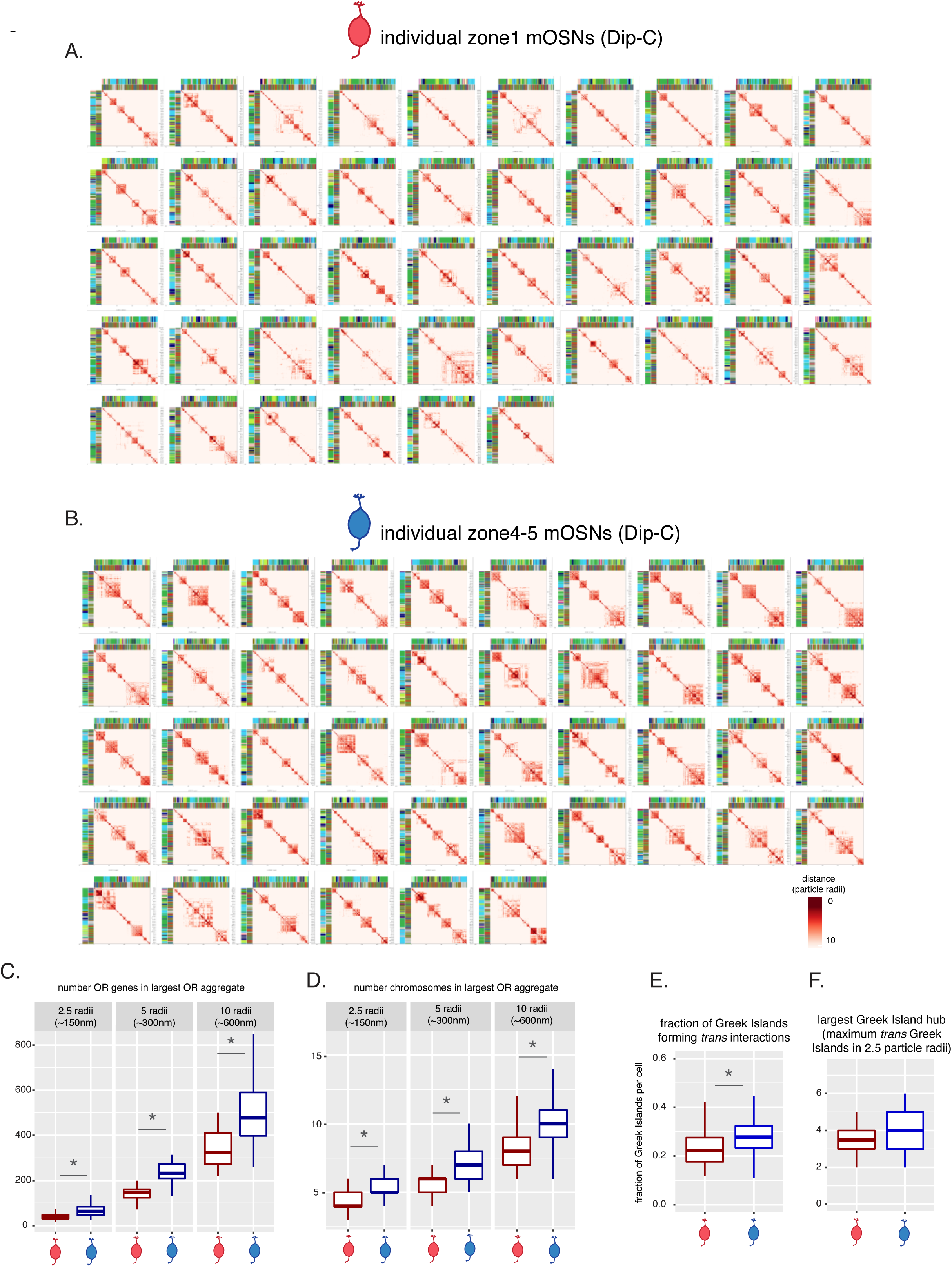
OR compartments are highly variable between single cells but show a consistent difference between cells of different zones. (A, B) Hierarchically clustered heatmaps of distances between zonal ORs (described in Figure 4) computed from 3D genome structures show OR genes within 10 particle radii (∼600nm) in each zone1 mOSN (A) and zone4/5 mOSN (B). (C,D) Analysis of all OR genes within 2.5, 5 and 10 particle radii (analogous to ∼150, 300 and 600nm) of one another shows that OR genes in zone 4-5 mOSNs are in proximity to a greater number of OR genes on different chromosomes (C) and form more complex compartments (a higher number of chromosomes within a given distance) (D). Asterisk denotes Wilcoxon rank sum test p-value <0.001). (E) On average a larger fraction of Greek Islands within zone 4-5 mOSNs is found near (within 2.5 particle radii) of Greek Islands from other chromosomes (mean of 27% in zone 4-5 cells, and 22% of zone 1 cells, Wilcoxon rank sum test p-value 0.003). (F) The size of the largest hub (defined as the maximum site of interchromosomal interactions between Greek Islands) --which ranges from 4-8 interchromosomal Greek Islands--is similar in zone 1 and zone 4-5 mOSNs.

**Supplementary Figure S5 (related to Figure 5):**
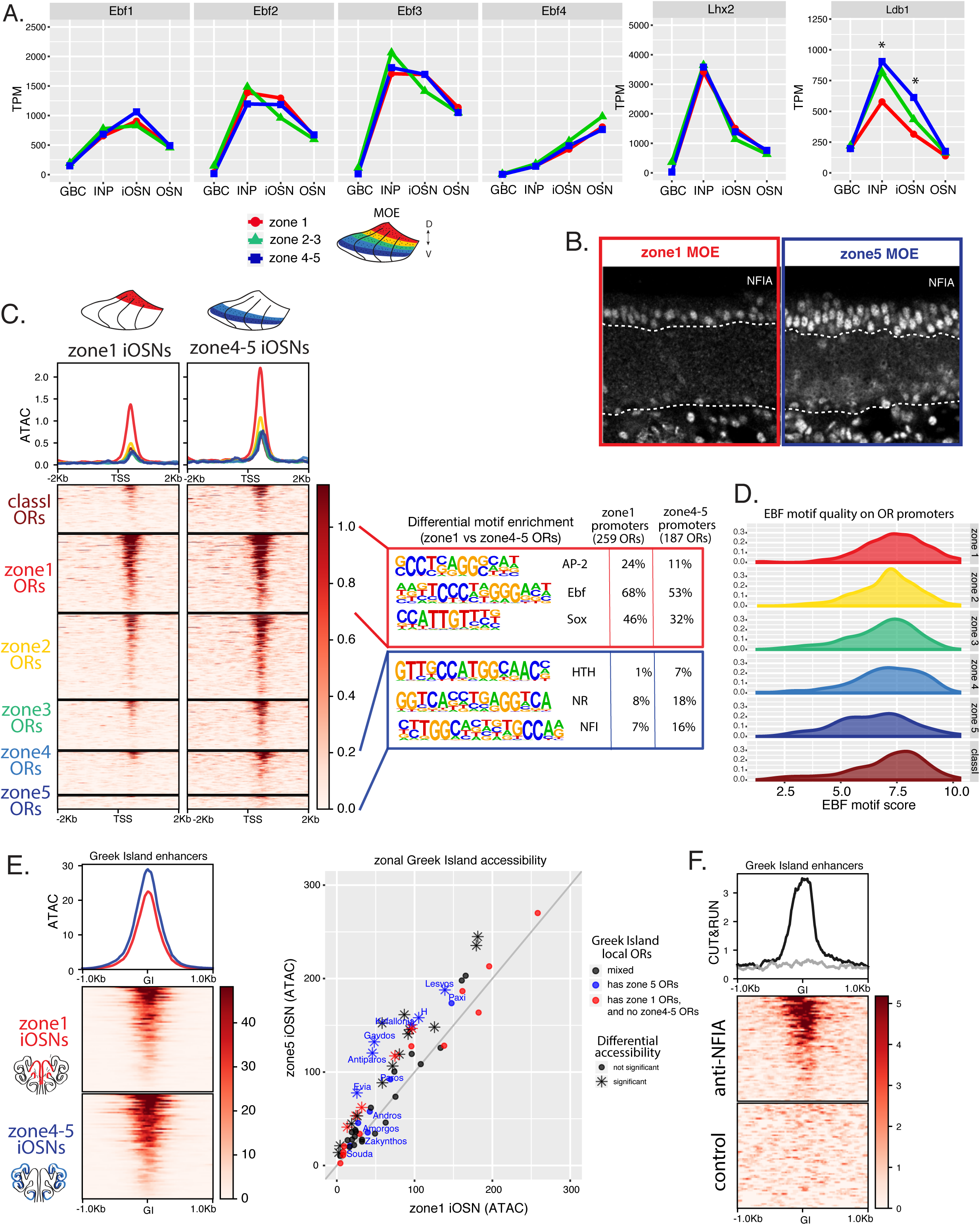
Increased expression of NFI factors in higher zones may compensate for their relatively weaker OR promoters. (A) Expression levels of core OSN lineage transcription factors EBF1-4, Lhx2 and Ldb1 at different stages of OSN development in zone1 cells (red), zone 2/3 cells (green) and zone 4/5 cells (blue). Out of these only Ldb1 is differentially upregulated in zone 4/5 in INP and iOSN cells (asterisk) with an adjusted p-value =0.007 and 2.08×10-7, respectively. (B) NFIA immunofluorescence staining in zones 1 and 5 MOE. Strong nuclear staining is only observed in zone 5 neuronal cells, which are outlined on the top and bottom with dashed lines. Staining is also observed in non-neuronal sustentacular cells (located above the dashed line) in both zones. (C) Heatmaps of omni-ATAC peaks in zone 1 and zone 4/5 iOSNs show increased coverage over OR promoters in zone 4/5 iOSNs. Heatmaps are centered on the TSS with 2kb flanking on either side and the plots above show the average signal per zonal category (colored by zonal annotation). The motifs shown were identified with HOMER to be differentially enriched on 259 zone 1 OR promoters relative to 187 zone 4/5 promoters and vice versa. Promoters are defined as 500bp upstream and downstream of the TSS. (D) Density plot of the best EBF motif found in each OR promoter separated by zone. (E) Heatmap of omni-ATAC signal on each of the 63 Greek Island enhancers in iOSNs, centered on the Greek Islands and showing a 1kb flanking region on either side (left). A plot of omni-ATAC signal on each Greek Island in zone 1 iOSNs vs zone 4/5 iOSNs (right). Greek Islands are colored based on the types of OR genes in their resident OR clusters (blue: clusters have 4/5 ORs; red: clusters have zone 1 ORs and no zone 5 ORs; black: all others). Significantly differentially accessible Greek Islands (p-value<0.05 determined with Diffbind) are shaped as stars. (F) Heatmap of NFIA CUT&RUN signal on Greek Islands shows enrichment of signal relative to a control antibody.

**Supplementary Figure S6 (related to Figure 6):**
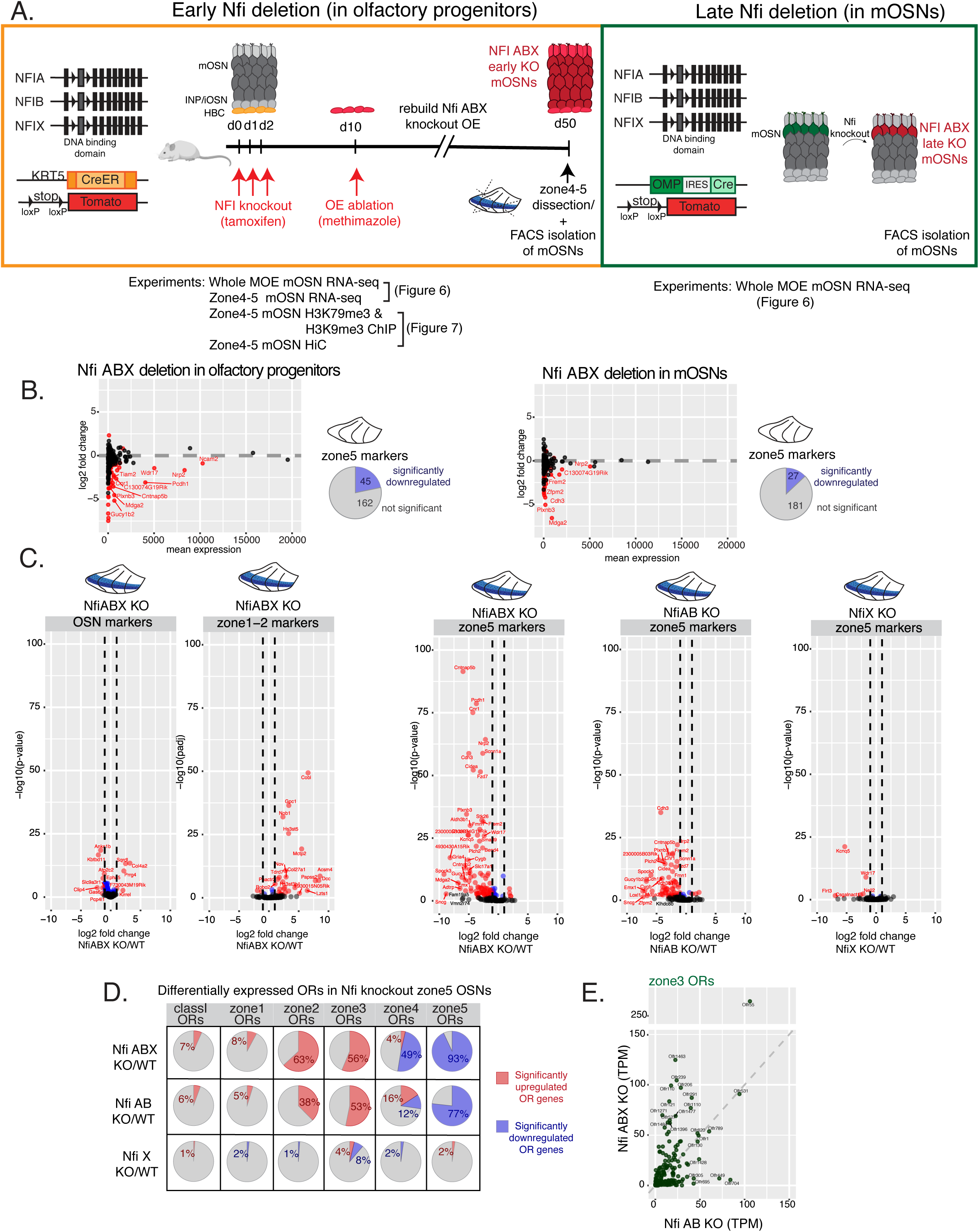
OE in Nfi ABX knockout zone5 undergoes a homeotic transition in their OR repertoire while maintaining their cellular identity. (A) Schematic showing the experimental and genetic strategy for deleting Nfi genes in progenitors (“early” deletion, left) and mOSNs (“late” deletion, right). (B) MA plot showing the effect of Nfi ABX “early” (left) and “late” (right) triple knockout on zone4/5 OSN marker expression (significantly differentially expressed genes are in red). Pie chart with quantification of significantly downregulated genes is shown. (C) Volcano plot showing expression of non-zonal mOSN markers, zone 1/2 mOSN markers and zone 4/5 mOSN markers in mOSNs isolated from NFI ABX triple, NFI AB double and NFIX zone4/5 MOE relative to wt. Only 13/200 non-zonal mature markers are significantly downregulated in NFI ABX triple knockout, compared to 90/207 zone 5 markers significantly downregulated and 29/138 zone1 markers that were significantly upregulated. Expression of zone 4/5 mOSN markers varied between the NFI knockout genotypes. Blue: significantly differentially expressed genes (p-value < 0.5), red: significantly differentially expressed genes with a > 2-fold change. (D) Differential expression analysis of all ORs in the different NFI knockout genotypes. Percentages of significantly upregulated (red) and downregulated (blue) ORs are shown. (E) Plot with expression of zone3 ORs in Nfi ABX triple and Nfi AB double knockout zone 4/5 shows a different set of zone3 ORs ectopically expressed in each genotype.

**Supplementary Figure S7 (related to Figure 7):**
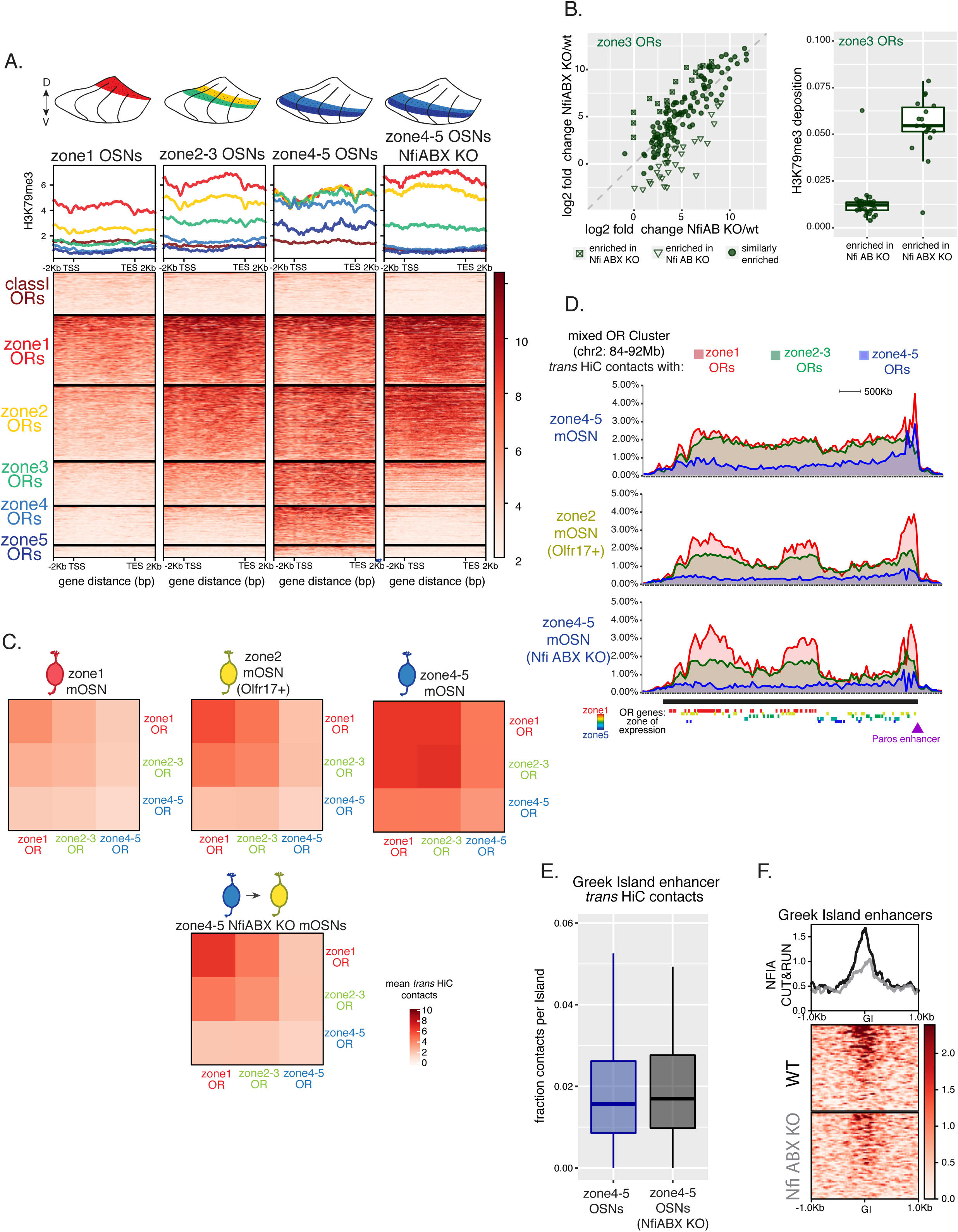
Nfi ABX knockout zone 4/5 cells closely resemble zone 2/3 cells in chromatin state, compartment formation and OR expression. (A) Heatmap with side by side comparison of H3K79me3 ChIP-seq signal over OR genes in mOSNs from zone 1, zone 2/3, zone 4/5 and Nfi ABX knockout zone 4/5 shows zone 4/5 knockout mOSNs have a chromatin state most similar to that of zone 2/3 mOSNs. (B) Identification of zone 3 OR genes that are differentially ectopically expressed in RNA-seq of zone 4/5 Nfi ABX triple knockout vs Nfi AB double knockout olfactory epithelium (left). Zone 3 OR genes that significantly more enriched in Nfi ABX triple knockout have a higher level of H3K79me3 in zone 4/5 NFI ABX knockout mOSNs. (C) Side by side heatmaps of average interchromosomal Hi-C contacts between OR genes of different zonal indexes show the nuclear OR interactome in zone 4/5 Nfi ABX knockout cells is most similar to that of zone 2 mOSN. (D) Density plot of trans Hi-C contacts made by zone1 (red), zone 2-3 (green) and zone 4-5 (blue) ORs with a large OR cluster on chromosome 2 shows a different pattern of interchromosomal interactions in each cell type. OR genes residing in the cluster are shown at the bottom (colored by their zonal index). Zone 4/5 mOSNs have a more level pattern of contacts, while both zone 2/3 mOSNs and zone 4/5 Nfi ABX knockout cells show preferential interactions with regions that are enriched for zone 1 ORs. Purple triangle marks the Paros enhancer. (E) In aggregate, there is no significant difference in intrachromosomal interaction between Greek Islands in zone 4/5 wt and NFI ABX triple knockout cells. (F) Nfia CUT&RUN signal over Greek Island enhancers in developing mOSNs isolated from the whole MOE of NFI ABX knockout and wt mice.

**Table 1.**
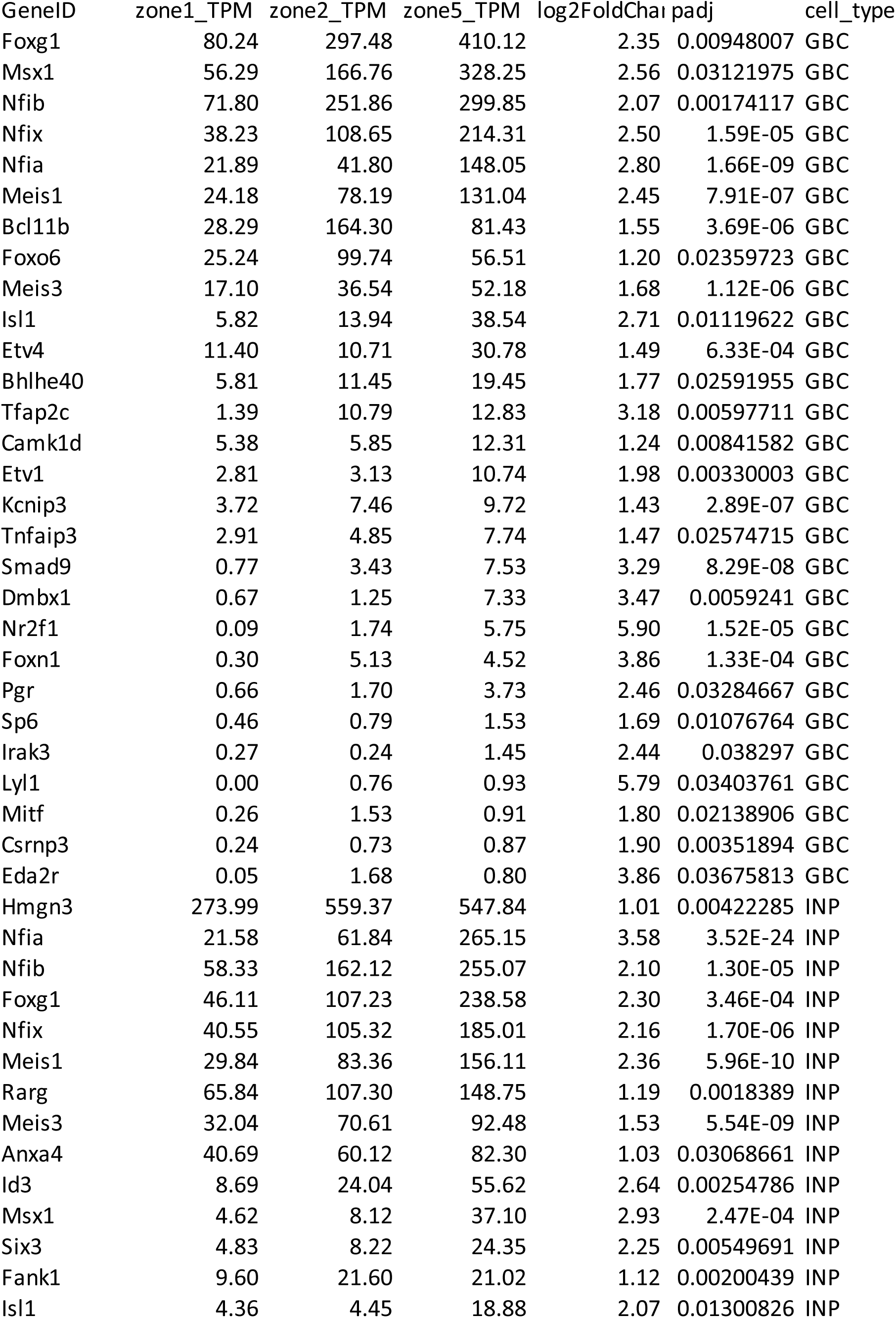
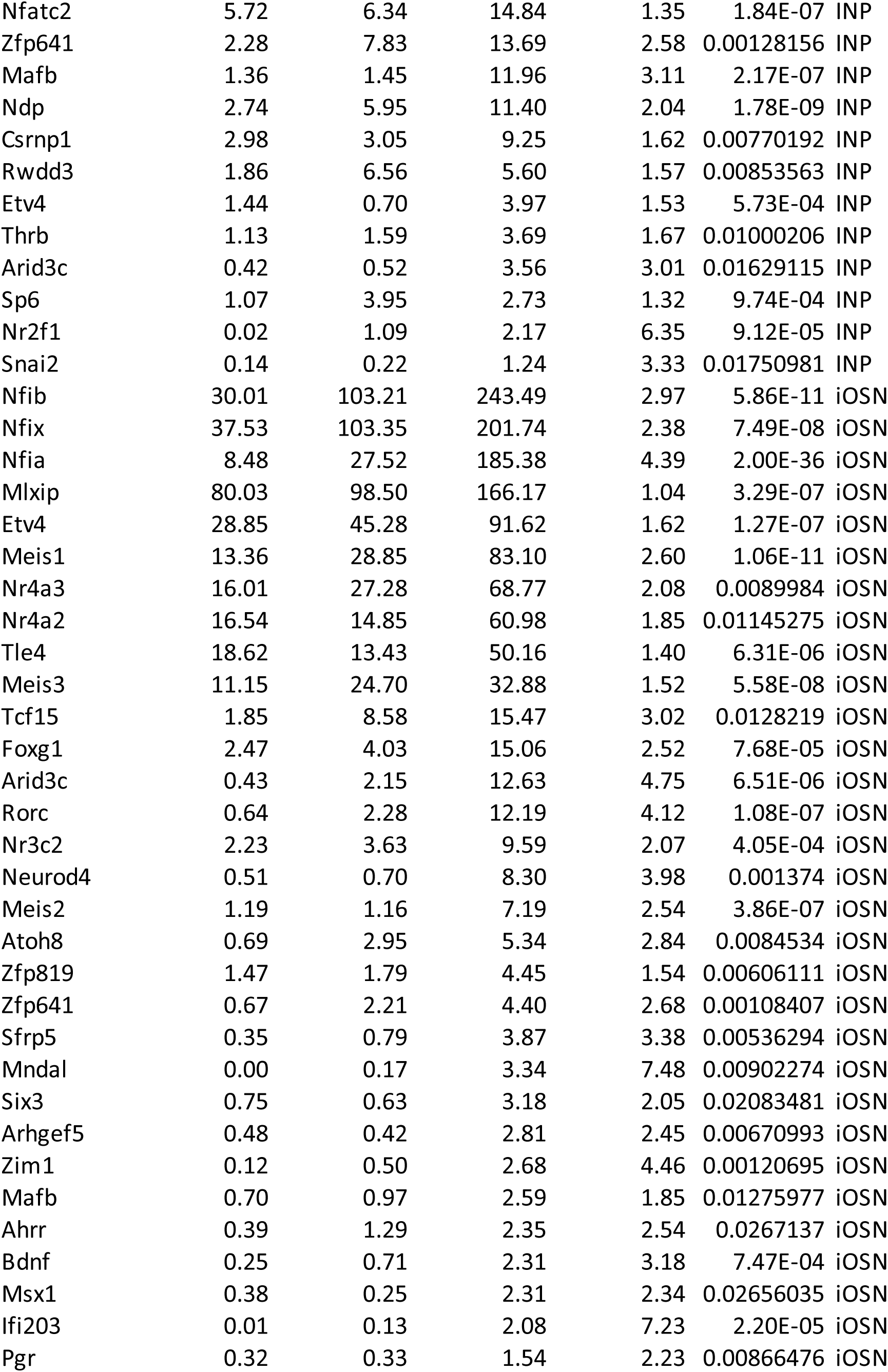
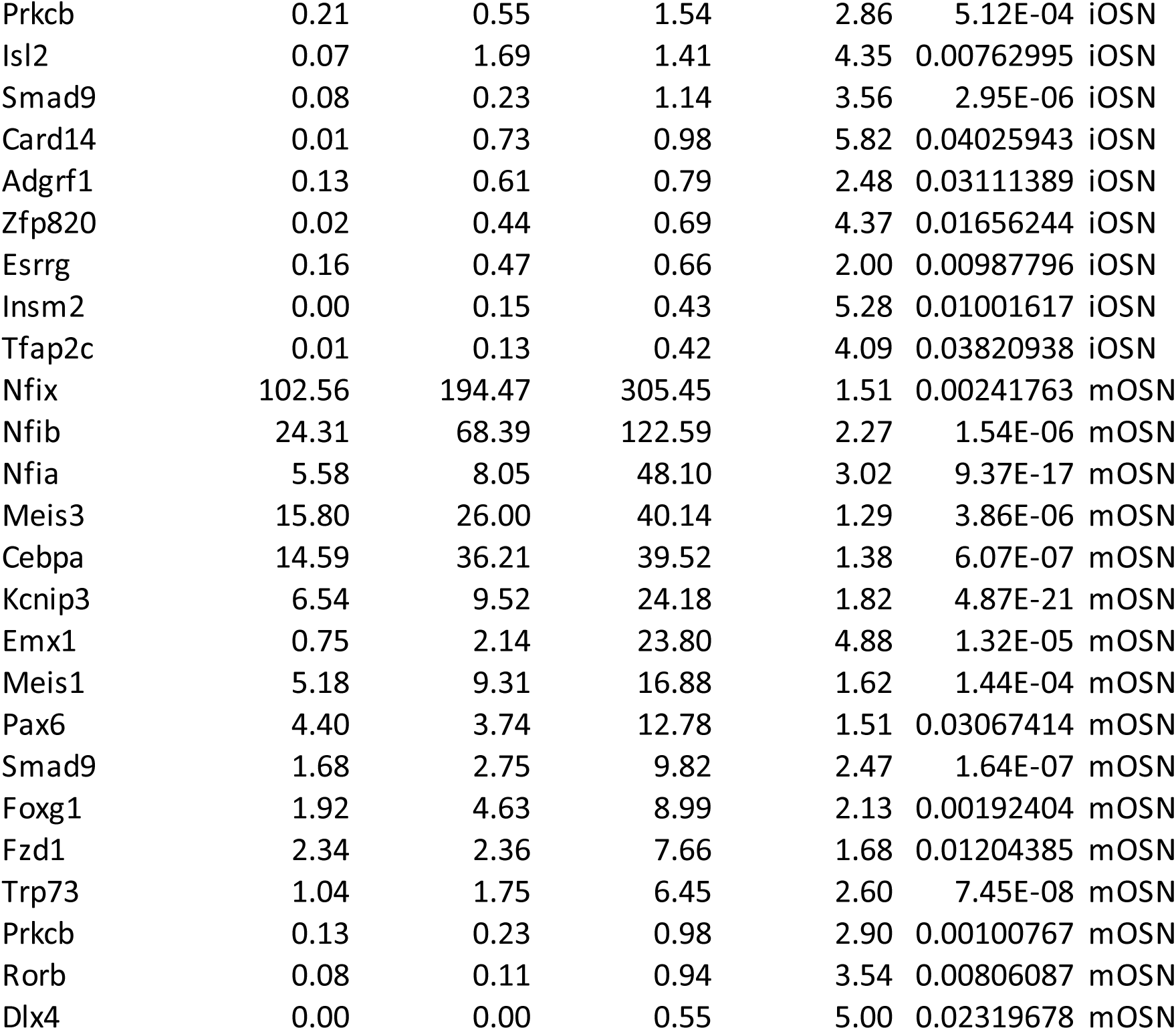
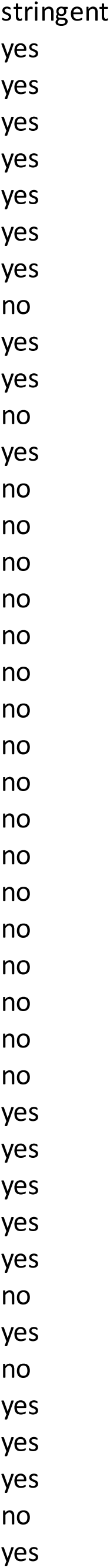
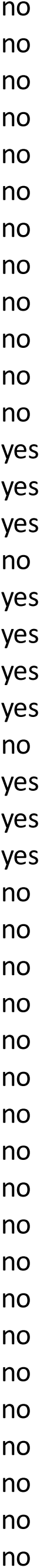
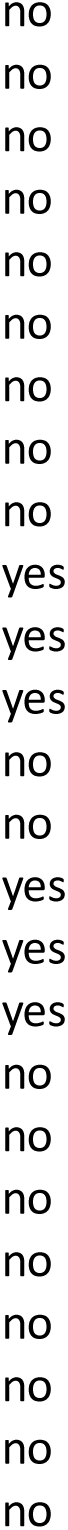
**(related to Figure 5):** Complete list of transcription factors differentially expressed between zones 1 and zones 4/5 at various stages of OSN differentiation. In the main Figure 5 we only included TFs with a 3 fold difference between the two zones but here we have a less stringent list including TFs with 2 fold differential expression.

**Table 2.**
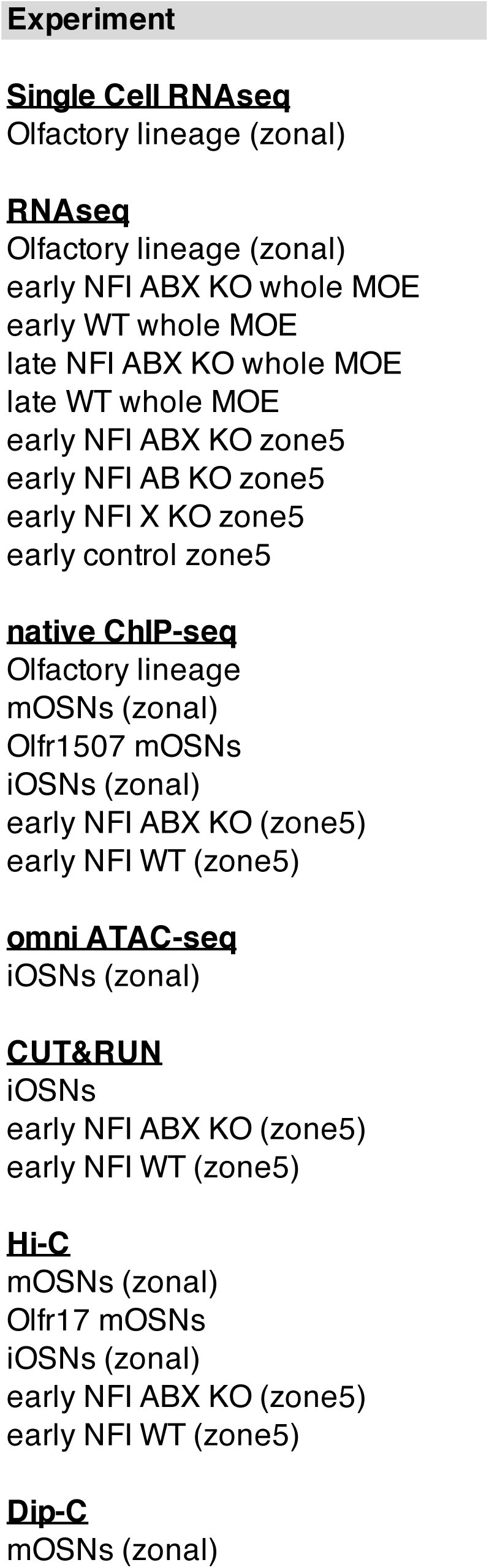
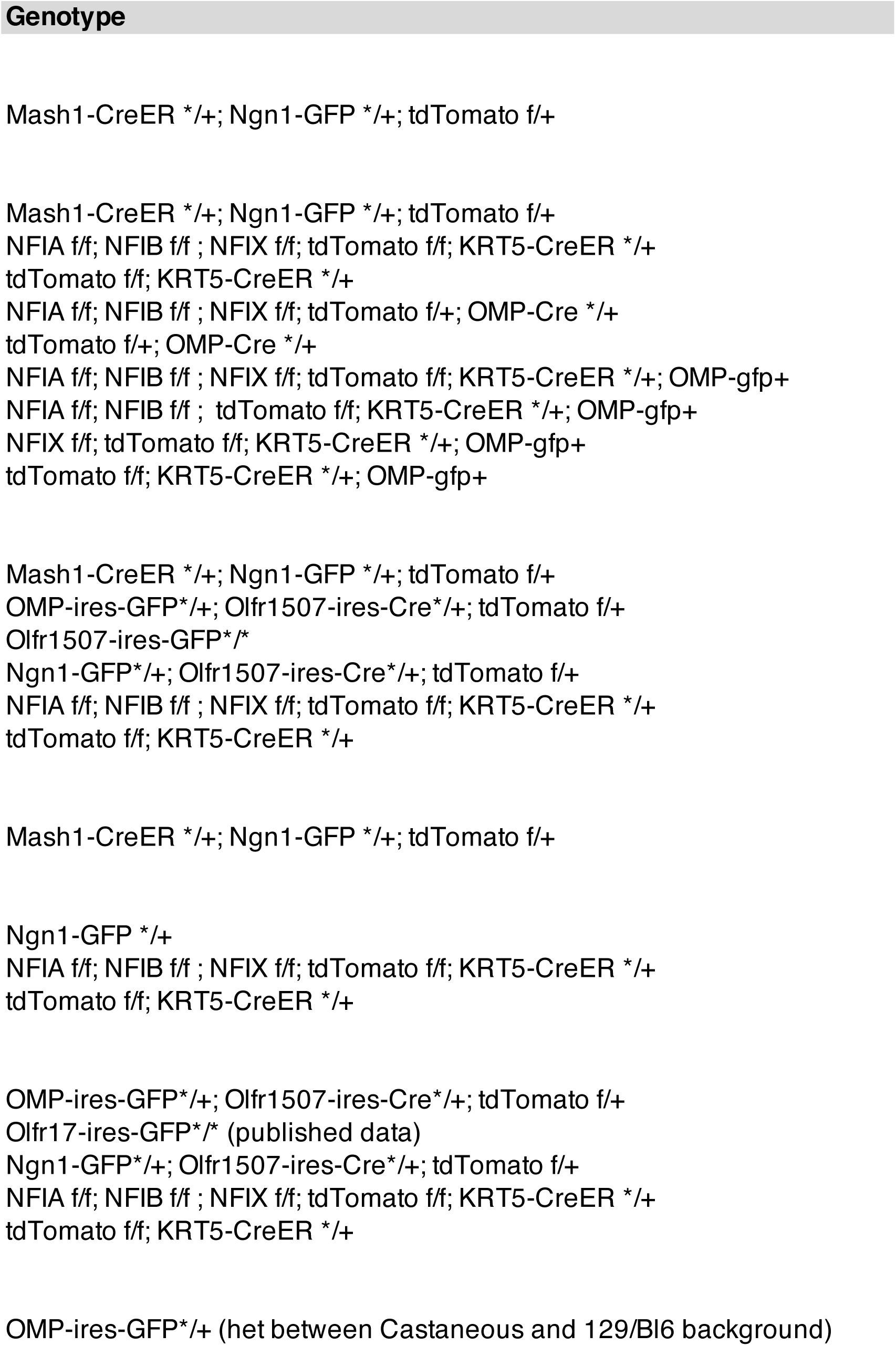
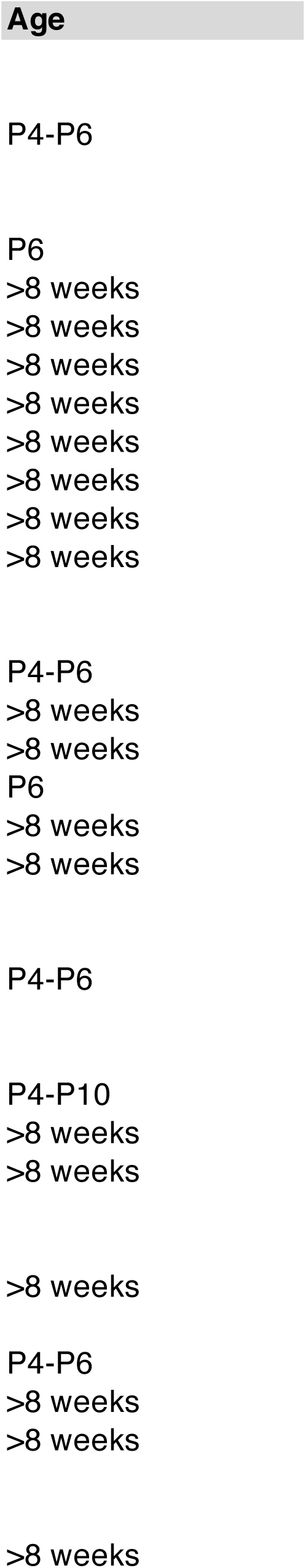
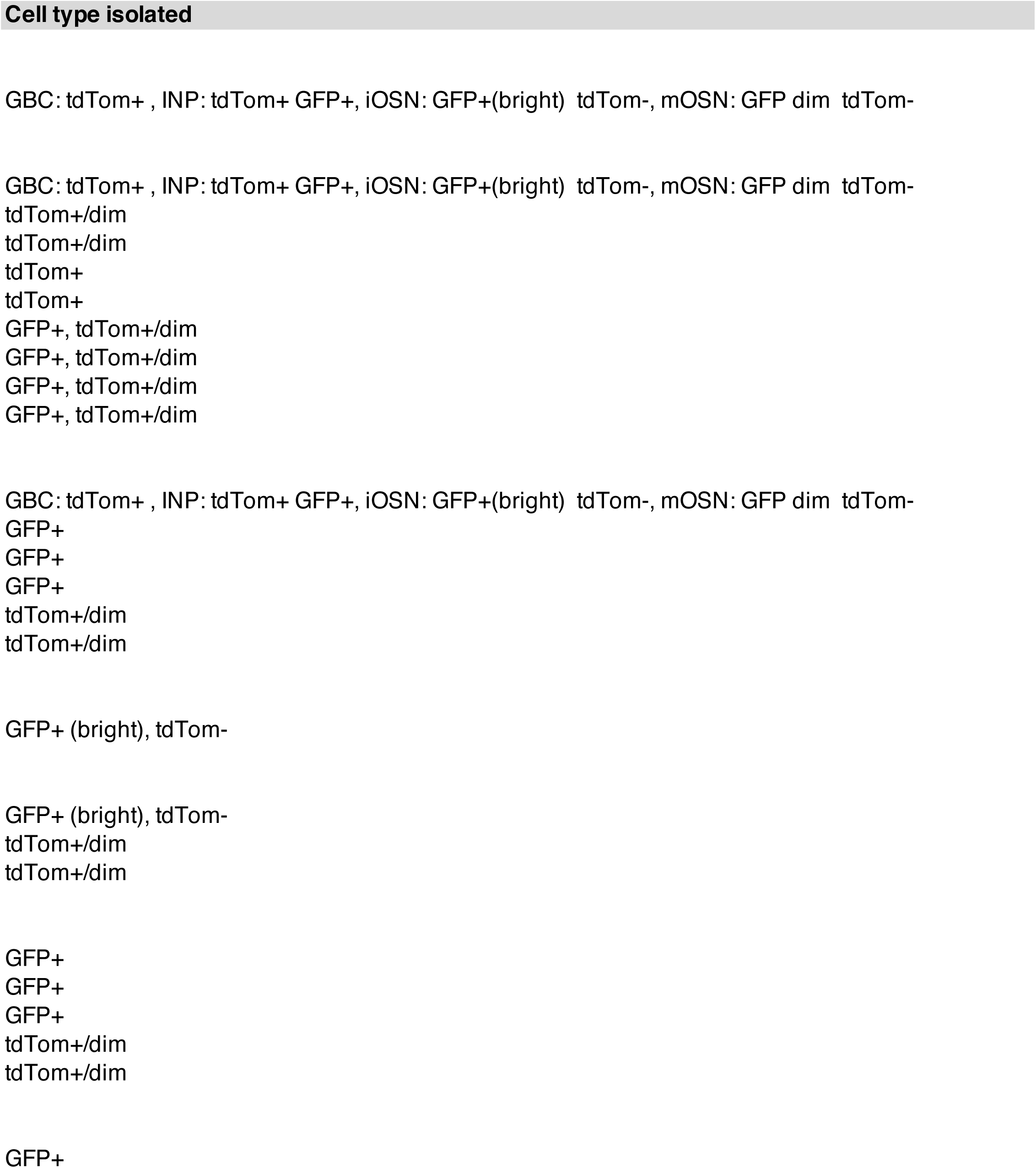
**(related to STAR methods):** Complete list of mouse genotypes analyzed in this manuscript.

